# Ultrastructural and Histological Cryopreservation of Mammalian Brains by Vitrification

**DOI:** 10.64898/2026.01.28.702375

**Authors:** Gregory M. Fahy, Ralf Spindler, Brian G. Wowk, Victor Vargas, Richard La, Bruce Thomson, Roberto Roa, Hugh Hixon, Steve Graber, Xian Ge, Adnan Sharif, Steven B. Harris, L. Stephen Coles

## Abstract

Studies of whole brain cryopreservation are rare but are potentially important for a variety of applications. It has been demonstrated that ultrastructure in whole rabbit and pig brains can be cryopreserved by vitrification (ice-free cryopreservation) after prior aldehyde fixation, but fixation limits the range of studies that can be done by neurobiologists, including studies that depend upon general molecular integrity, signal transduction, macromolecular synthesis, and other physiological processes. We now show that whole brain ultrastructure can be preserved by vitrification without prior aldehyde fixation. Rabbit brain perfusion with the M22 vitrification solution followed by vitrification, warming, and fixation showed an absence of visible ice damage and overall structural preservation, but osmotic brain shrinkage sufficient to distort and obscure neuroanatomical detail. Neuroanatomical preservation in the presence of M22 was also investigated in human cerebral cortical biopsies taken after whole brain perfusion with M22. These biopsies did not form ice upon cooling or warming, and high power electron microscopy showed dehydrated and electron-dense but predominantly intact cells, neuropil, and synapses with no signs of ice crystal damage, and partial dilution of these samples restored normal cortical pyramidal cell shapes. To further evaluate ultrastructural preservation within the severely dehydrated brain, rabbit brains were perfused with M22 and then partially washed free of M22 before fixation. Perfusion dilution of the brain to 3-5M M22 resulted in brain re-expansion and the re-appearance of well-defined neuroanatomical features, but rehydration of the brain to 1M M22 resulted in ultrastructural damage suggestive of preventable osmotic injury caused by incomplete removal of M22. We conclude that both animal and human brains can be cryopreserved by vitrification with predominant retention of ultrastructural integrity without the need for prior aldehyde fixation. This observation has direct relevance to the feasibility of human cryopreservation, for which direct evidence has been lacking until this report. It also provides a starting point for perfecting brain cryopreservation, which may be necessary for lengthy space travel and could allow future medical time travel.

## Introduction

Cryopreservation of the mammalian brain has been studied or proposed as a method of analyzing the metabolism of single cells within the intact brain using the 2-deoxyglucose method [[1]], studying the tolerance of the brain to ischemia [[2]], studying the resistance of whole mammals to freezing [[3, 4]], mapping the connectome and facilitating other neurobiological techniques [[5]], advancing basic cryobiological research [[6]], improving brain banking [[7]], and developing medical applications [[8–11]]. Suda et al. reported that a cat brain frozen to-20°C with 15% v/v glycerol and stored for 5 days was able to generate electroencephalographic activity after thawing that was similar to the activity obtained from the same brain prior to freezing [[2, 6]]. Follow-up studies by the same authors showed that glycerol was superior to dimethyl sulfoxide at-20°C and that storage with glycerol at-20°C gave “well synchronized discharges of Purkinje cells,” spontaneous electrical activity in the thalamus and cerebellum, and a brief return of weak cortical electrical activity even after up to 7.25 years of storage, albeit with strong reduction of cell density and severe hemorrhage after this extreme period of storage [[6]]. Although unit activity could be recovered after freezing to-60°C, EEG activity could not (Suda et al., unpublished observations), and all electrical activity was abolished after cooling to-90°C [[6]]. In the absence of glycerol, Lovelock and Smith observed that whole hamsters could be frozen to a colonic temperature of-1°C with subsequent survival of the animals after thawing. This resulted in brain temperatures of-0.75 to-0.9°C, which corresponded to the freezing of 53-63% of the water in the brain based on both calculations and calorimetry [[3]]. Breathing and shivering were observed even after up to 70% of brain water had frozen [[4]]. However, in the Suda experiments, “pial oozing” was noted during blood reperfusion [[2]], which was later shown to be associated with fissures in the tissue ([[6]]; Suda, unpublished observations), which in turn are likely due to the prior formation of large ice cavities in the brain tissue (Fig. 1). Consequently, current information suggests that vitrification [[13]], or “ice-free cryopreservation”, which can prevent ice cavities in whole brains [[5, 9]], is a more promising method for brain cryopreservation than is freezing, and this possibility is supported by the preservation of excellent viability and ultrastructure [[14]] as well as normal electrical responsiveness of adult rat [[15]] and rabbit [[16]] hippocampal slices after vitrification, but not after freezing [[14]]. Consequently, vitrification is the approach that is further investigated here.

**Fig 1.**
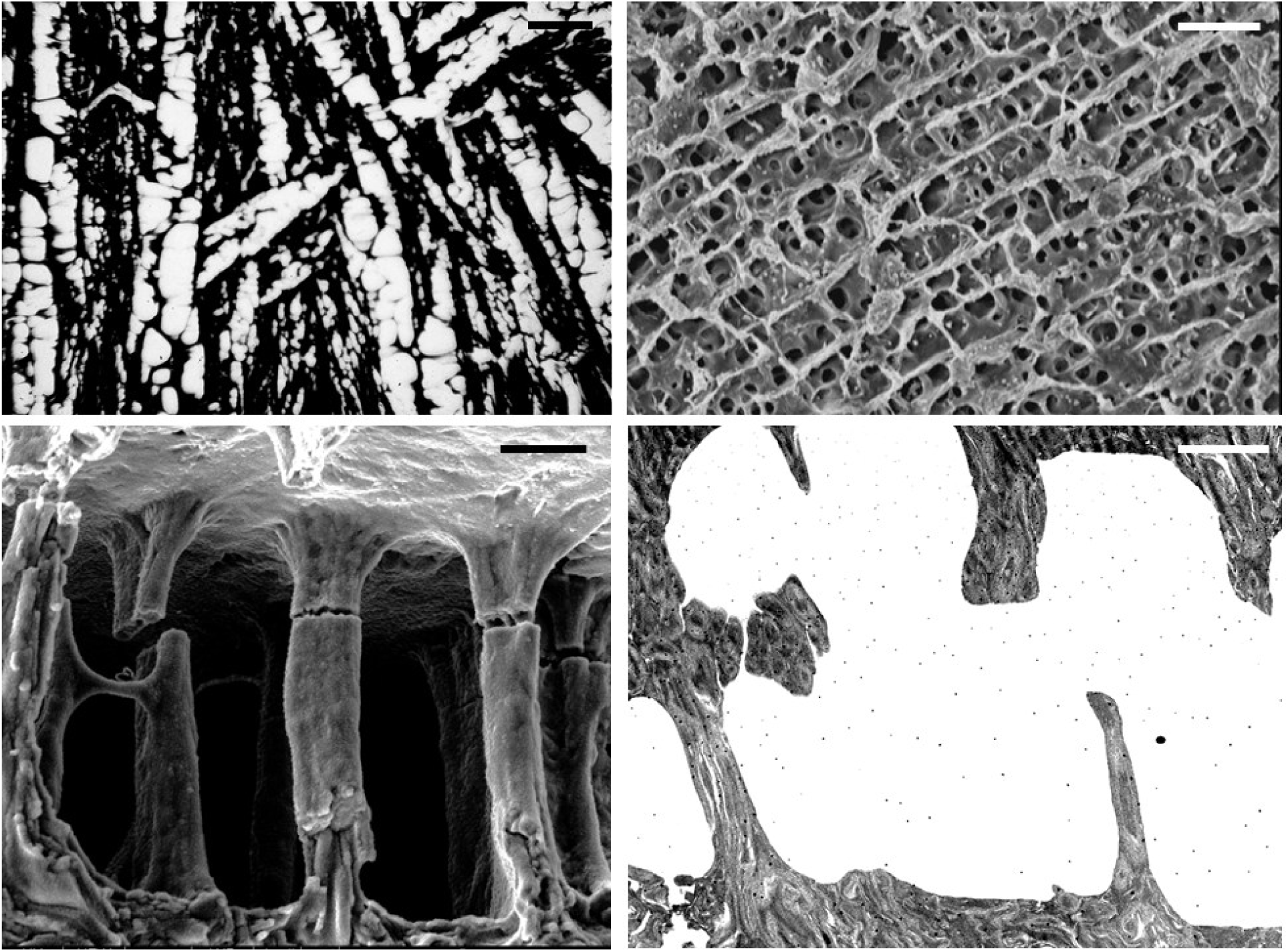
Distortion of gross rabbit brain histological morphology and ultrastructure by slow freezing with 3.72M glycerol. Light microscopic (**upper left),** scanning electron microscopic (upper right and lower left), and transmission electron microscopic (**lower right)** images of cerebral cortex removed from a rabbit brain perfused with 3.72M glycerol as described elsewhere [[1]], slowly frozen to-79°C, and freeze-substituted at this temperature [[12]]. Gross distortion of brain structure by large ice dendrites is evident (upper images), although the ice cavities remaining after freeze-substitution have smooth walls, and structural detail within the compacted tissue is visible (lower images). Scale bars indicate approximately 30 microns in the upper images and 2 microns in the lower images.

A significant obstacle to brain cryopreservation by either freezing or vitrification arises from the osmotic effects of cryoprotective agents, but these effects are particularly pronounced when attempting vitrification. For example, the M22 vitrification solution has an approximate melting point of-55°C [[17]], which implies an osmotic concentration of about 55/1.86 = 29.6 osmolal, which is 100 times more concentrated than plasma or cerebrospinal fluid. The inability of perfused cryoprotective agents to pass through the blood-brain barrier (BBB) and into brain cells and processes as fast as water can move by osmosis in the opposite direction results in significant osmotic brain dehydration and shrinkage (Fig. 2) [[1, 5, 9]]), and this shrinkage distorts and obscures normal histological [[1]] and ultrastructural [[9]] features. What is not known is whether this distortion is reversible or is sufficient to cause irreversible structural changes to the brain that may potentially lead to a loss of biological information.

**Fig. 2.**
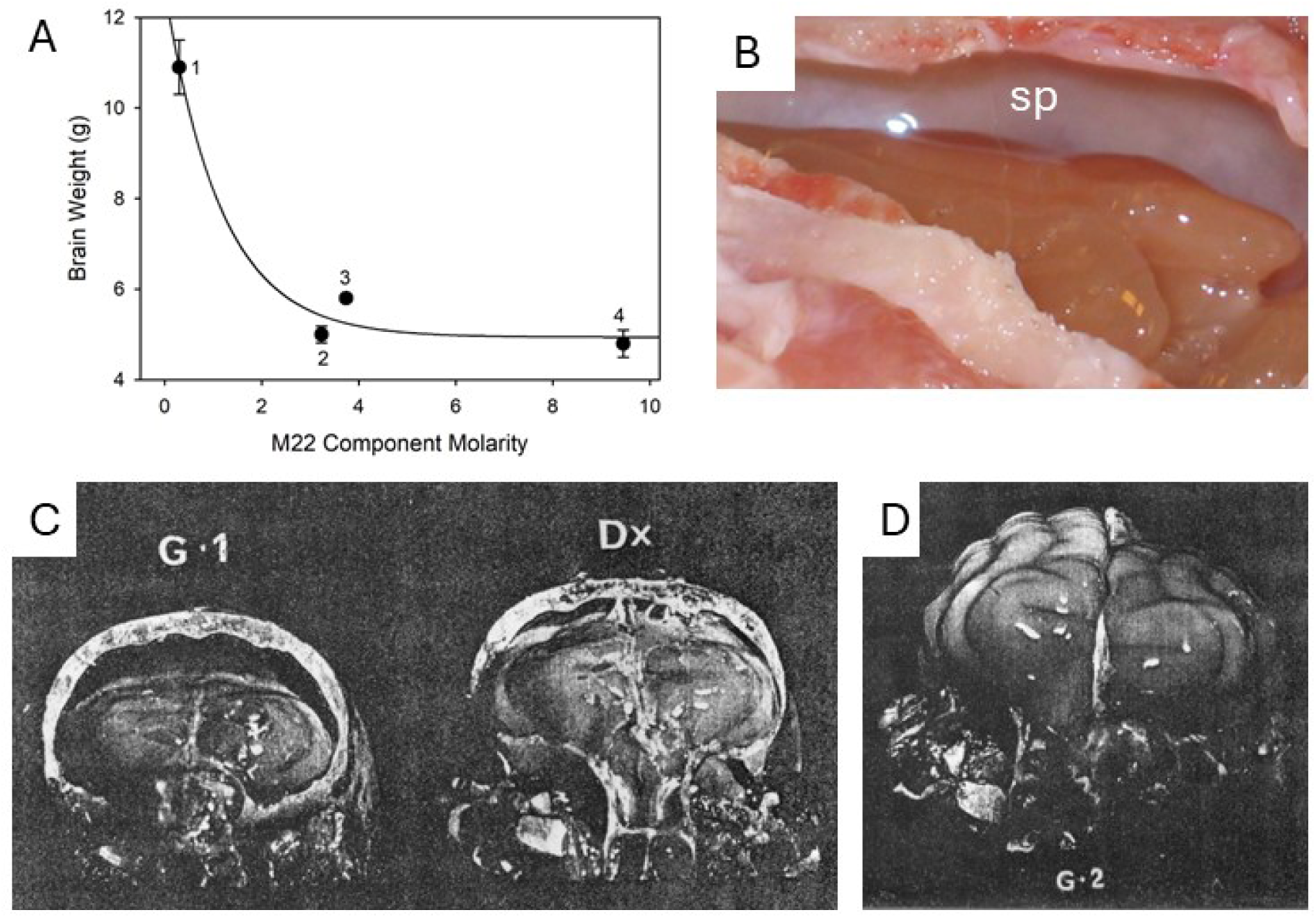
Typical examples of brain shrinkage induced by cryoprotectant perfusion and brain swelling upon cryoprotectant washout. **A**. Effect of cryoprotectant concentration on brain weight. 1 = no cryoprotectant (n=3). 2 = the ethylene glycol, 3-methoxy-1,2-propanediol, and polymer (X-1000, Z-1000, and PVP) components of M22 only (n=3). 3 = 2 plus the N-methylformamide component of M22 (n=3). 4 = the complete M22 composition (effective total molarity, 9.45M) (n=6). Data are in general accord with Boyle-van ‘t Hoff behavior [[18]] (plot not shown). **B**. Rabbit brain shrinkage by perfusion with M22, accompanied by formation of a shrinkage space (sp) between the skull and the amber-colored, shrunken brain. **C**. Similar shrinkage of the cat brain after perfusion with glycerol (G1) but not with Hanks-dextran (Dx). **D**. Return of the volume of a glycerolized cat brain to normal after reperfusion with Hanks-dextran (G2; in this image, the hemisphere seen on the right was removed following glycerolization and remains shrunken, whereas the hemisphere seen on the left was reperfused, after which it was returned to the skull for size comparison. C and D: unpublished images of Suda, Kito, and Adachi, published by permission of Isamu Suda.

Some shrinkage of the brain and of brain cells is known to be tolerable. Brain cells are believed to shrink mildly during sleep to facilitate removal of waste products from the brain via the glymphatic system [[19, 20]]. In addition, mild brain shrinkage has been studied in vivo [[21]] and used to treat neurological problems [[22]] and is likely present during attempts to open the blood-brain barrier by osmotic means for intracerebral drug delivery [[23]], with acceptable outcomes. The freezing of brain tissue samples also causes major shrinkage of neurons and neuronal processes but is compatible with strong recovery of neuronal functions upon thawing [[24–26]]. Moreover, in Lovelock and Smith’s experiments, the formation of ice crystals within the intact brain undoubtedly induced major shrinkage of cerebral neurons and their associated processes [[27]], but the hamsters apparently recovered neurologically after this insult, and similar brain shrinkage also occurs in freeze-tolerant vertebrates under natural conditions, with no harmful sequelae [[28, 29]]. The most direct evidence comes from Suda’s studies [[2, 6]], which showed that despite major brain shrinkage induced by glycerol perfusion followed by additional shrinkage caused by freezing to-20°C (which can be estimated [[30]] to result in conversion of approximately 61% of brain liquid volume to ice under equilibrium conditions) and brain re-expansion to its original or greater than original size after perfusion with a glycerol-free solution (Fig. 2C and 2D), spontaneous restoration of electrical activity was still possible. Nevertheless, Suda did not investigate the ultrastructural state of glycerolized or frozen-thawed brains, and he did not establish the limits of safe brain shrinkage. Therefore, given the very high concentrations of cryoprotectants required for vitrification [[13]], it is possible that the preparation of brains for vitrification could exceed the still-undefined safe limits of shrinkage of the whole brain, and previous descriptions of brain structure after cryoprotectant perfusion [[1, 9]] were not sufficiently detailed to answer this question.

We therefore addressed this question by examining brains at higher magnification in the presence of extreme dehydration after vitrification and rewarming using light microscopy and low and high magnification electron microscopy. We also employed graded washout of cryoprotectant to partially rehydrate the brain prior to fixation and examination. The results indicate that both animal and human brains can retain overall ultrastructural integrity after vitrification and rewarming without prior aldehyde fixation.

## Results

### Brain Ultrastructure and Histology after Vitrification and Rewarming

**Fig. 3** Illustrates the typical overall features of the rabbit brain after perfusion with M22 for one hour, vitrification, rewarming, and perfusion-fixation with M22 fixative, as visualized by low and higher power scanning electron microscopy. The black box in **Fig. 3A** designates regions of the hippocampus and the overlying cerebral cortex that were selected for more detailed TEM and STEM observations in similarly-treated brains as described in subsequent Fig.s. The small white box and the large white box in Fig. 3A denote regions displayed at higher magnification in Fig.s 3B and 3C, respectively. Low power imaging (**Fig. 3A**) shows an absence, over macroscopic distances, of any evidence of structural disruption from ice crystals, in contrast to the phenomena described in **Fig. 1**. Overall brain structures appear to be intact, although larger hippocampal blood vessels are visibly dilated. Higher power imaging of the CA1 cell band of the hippocampus (**B**) reveals, consistent with the macroscopic shrinkage of the whole brain (**Fig. 2**), very dehydrated and distorted but structurally intact CA1 cells and their intact and well-aligned apical axons, with extracellular shrinkage spaces between smooth and continuous cellular surfaces (heavy arrows). Capillaries appear to be intact (thinner arrows). Intact capillaries are visible throughout the brain, and are further highlighted in **Fig.s 3C and 3D**. As displayed in **Fig. 3C**, some hippocampal arterioles show separation from otherwise intact surrounding neuropil (upper white arrow) while others show retraction of the arteriole proper away from its own outer layer, which remains adjacent to the neuropil (lower white arrow). The gaps in either case are consistent with possible osmotic and hydrostatic distension during cryoprotectant loading followed by hydrostatic contraction after perfusion cessation. However, capillaries are not affected by such phenomena (black arrow). **Fig. 3D** shows a higher magnification image of the cerebral cortex. The capillaries in the cortex, like those in the hippocampus, appear universally open and intact (arrows). Double arrows show where one particular capillary enters and exists the plane of the section, with a smooth and intact capillary wall between these two points. In general, the tissue is intact, with unremarkable texturing.

**Fig. 3.**
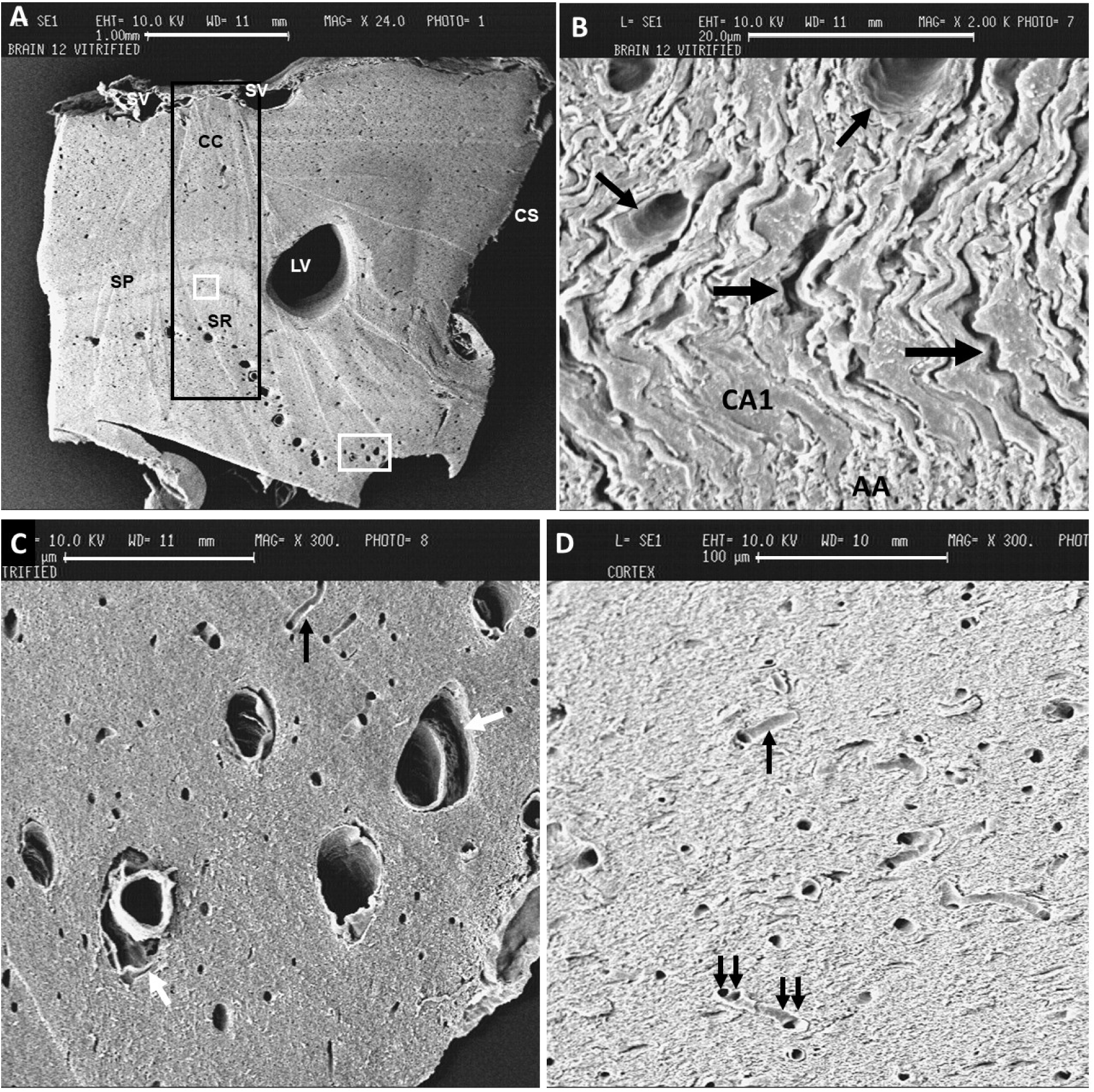
Scanning electron microscope images of the dorsomedial area of one hemisphere of the rabbit brain after vitrification, rewarming, fixation in the presence of the cryoprotectant, cryoprotectant elution, and hand-cutting with a Stadie-Riggs microtome blade, showing the general structural features of the vitrified and rewarmed brain. **A.** Overall integrity and structure of the brain. The black rectangle denotes the regions of cortex and hippocampus whose ultrastructural details are described in more detail in subsequent Fig.s. Tissue collection was standardized by sampling the hippocampus at the most dorsal region of the CA1 cell band and the cerebral cortex directly above it. CC, cerebral cortex; SP, stratum pyramidale; SR, stratum radiatum; LV, lateral ventricle; CS, central sulcus. Surface veins (SV) are adjacent to the cerebral cortex. The upper white rectangle and the lower white rectangle denote the regions displayed at higher magnifications in **B** and **C**, respectively. Scale bar: 1 mm. **B.** CA1 band of the hippocampus (CA1) and its underlying apical axon field (AA) descending into the SR. Arrows indicate intact capillaries. Heavy arrows indicate intercellular shrinkage spaces. Scale bar: 20 microns. **C.** Hippocampal arterioles, intact except for occasional delamination (lower white arrow) or smooth gaps between the arterioles and the surrounding neuropil (upper white arrow). Capillaries are smooth, intact, and adjacent to surrounding neuropil (black arrow). Scale bar: 100 microns. **D**. General appearance of the cerebral cortex, showing smooth, intact capillary walls (arrow) with no separation between capillaries and surrounding neuropil. Capillary continuity between at points of capillary entry into and exit from the plane of the section (double arrows) is evident. The texture of the neuropil between capillaries shows relief from cut neuronal membranes. Modified from [[9]]. Scale bar, 100 microns.

**Fig. 4** describes a number of salient features observed in an exemplary rabbit brain that was perfused with M22 for 60 min, vitrified, rewarmed, and fixed with M22 fixative. Evidence shown in Fig. 3 for a general absence of ice-induced tissue cavities is confirmed in all brain regions examined by TEM. Neuronal shrinkage and shape distortion are apparent in both the cortex (**A, B**) and hippocampus (**D-F**), obscuring intracellular detail, although cell nuclei can be seen at intermediate magnifications (**B, D-F**). At low magnification (**A**), tissue dehydration can be appreciated by the overall increased density of the tissue and by frank cellular shrinkage (arrows), without apparent loss of cellular contents, and extracellular shrinkage spaces are visible in both cortex (**A-C**) and hippocampus (**D-F**). At higher magnifications, intact mitochondria (m), synapses (short arrow), and cellular and neuronal process membranes can be observed (**K**). As also seen by SEM, separation of some arterioles from the surrounding neuropil is present (**C**), but capillaries are consistently intact and remain in normal contact with surrounding parenchyma (**A, E, G**). Small (**G, K**) myelinated fibers and their myelin sheaths remain generally intact, but larger myelinated fibers can show partial separation of myelin layers (**J**) and shrinkage spaces between intact axons and their myelin sheaths and between their myelin sheaths and the surrounding neuropil (**H-J**). Potentially incomplete osmication may occur in some cells (D), potentially due to inaccessibility of reactive tissue sites due to extensive tissue dehydration. Either incomplete osmication or localized intracellular protein precipitation can also be observed at higher magnification (long arrow in **K**). However, there is no suggestion of interruption of neuronal processes or their synaptic connections. The overall appearance is one of preservation of molecular, structural, and connectomic detail, with some changes whose potential reversibility requires rehydration to evaluate.

**Fig. 4.**
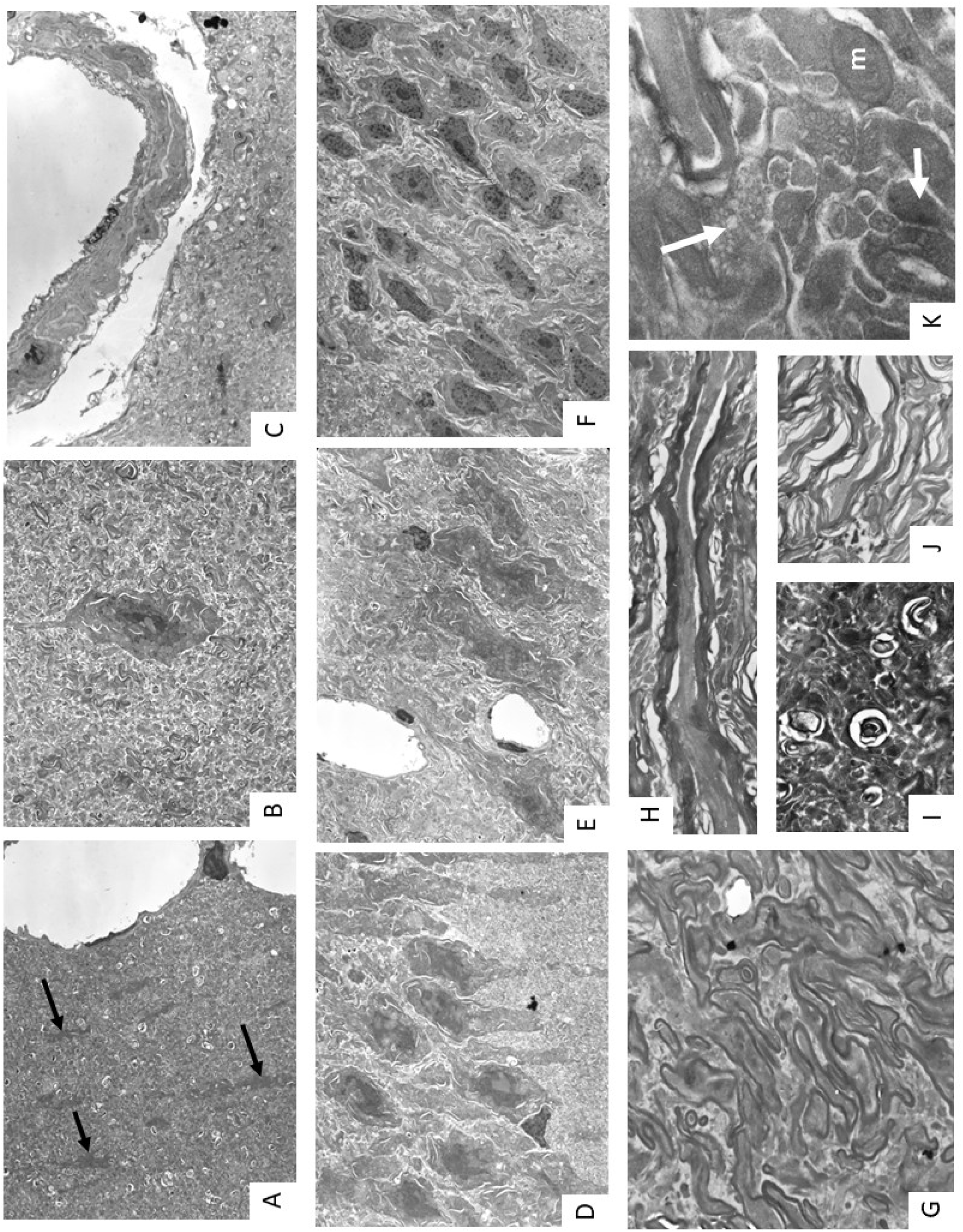
Typical appearance of the M22-perfused, vitrified, rewarmed, and immediately perfusion-fixed rabbit brain. **A**. Relatively low-power image showing shrunken cerebral cortical cells (arrows) embedded in dehydrated but intact neuropil with intact capillaries nearby. There is no apparent loss of ground substance. **B.** Higher power image showing a shrunken and distorted but evidently intact pyramidal cell surrounded by neuropil. White shrinkage spaces are visible within the neuropil. **C.** Separation of a larger blood vessel from the surrounding neuropil. Despite separation, the blood vessel and the surrounding neuropil are largely intact. **D**. Shrunken but well-aligned CA-1 cells, with minor extracellular and intracellular [[31]] shrinkage spaces typical of dehydrated cells; osmication appears incomplete in some areas, possibly due to dehydration. **E**. Intact capillaries in another CA-1 region. **F.** Dehydrated but intact dentate gyrus granular cells, showing small paracellular shrinkage spaces. **G**. Intact small para-hippocampal myelinated axons with well-apposed axoplasm and myelin. **H**. Longitudinal section through a larger hippocampal myelinated axon, showing shrinkage of the axon within the myelin sheath and shrinkage spaces external to the sheath. **I**. Hippocampal axons in cross section, showing periaxonal shrinkage spaces in the mossy fiber field. **J**. Shrunken para-hippocampal myelinated fibers showing partial separation of intact myelin layers. **K**. High magnification image showing an intact myelinated fiber and intact, well-defined unmyelinated cell processes but signs of coagulated or precipitated cytoplasm (upper arrow). An intact synapse (lower arrow) and an intact mitochondrion (m) are also visible. Experiment RM2201.

For comparison, we evaluated the structural state of cerebral cortical biopsies that were taken from an M22-perfused human brain and placed directly into liquid nitrogen for later examination. The biopsies were divided into smaller samples under liquid nitrogen and then warmed by placement in M22 fixative or in diluted M22 (see below) followed by diluted M22 fixative. As shown in **Fig. 5A**, the samples appeared amber-colored prior to (top) and following (bottom) warming, which is typical of tissue saturated with M22, and they did not exhibit whitening suggestive of ice formation during warming, implying that the tissue had vitrified on cooling and had remained ice-free on warming. These visual impressions were confirmed by differential scanning calorimetry (DSC). Of three samples tested by DSC, none formed ice during cooling at 1°C/min to-90°C, below which ice growth is negligible [[32]] (see Supplementary Materials). Absence of ice at this cooling rate is consistent with the vitrifiability of the human brain even at rates achievable by normal external cooling. Further, no DSC sample formed ice even under conditions formerly shown to nucleate and grow ice in 90% M22 [[32]] (see Supplementary Materials), implying that the effective M22 concentration in the cerebral cortex was over 90% of that in full M22. In support, direct HPLC measurements of tissue cryoprotectant concentrations combined with tissue dehydration are consistent with strong tissue resistance to ice formation (see Supplemental Materials).

**Fig. 5.**
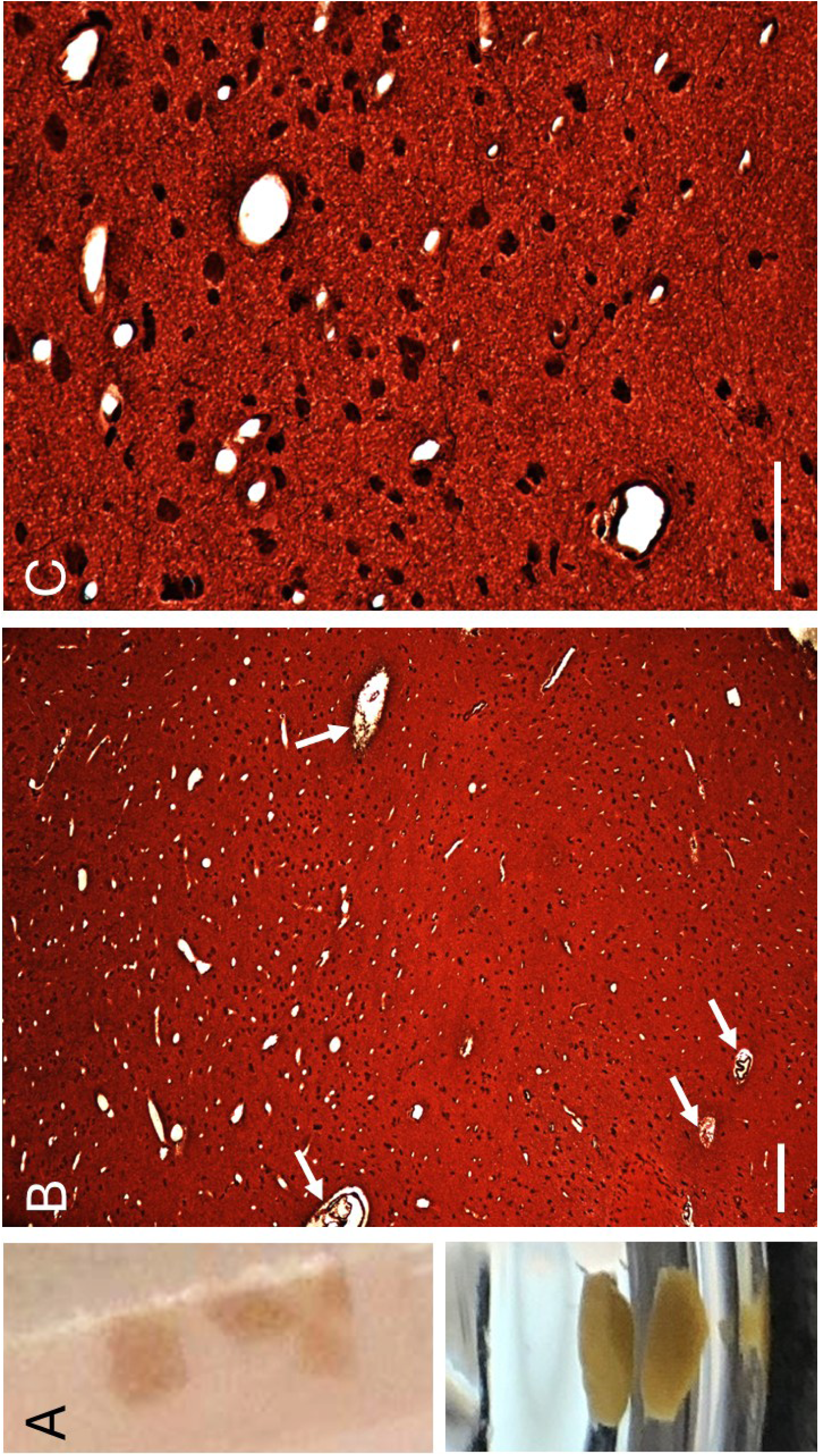
Pre-and post-warming state and histological integrity of human cerebral cortical tissue following whole brain perfusion with M22 and biopsy storage in liquid nitrogen for 4 years. **A.** Typical amber-colored appearance of biopsies in a Nunc tube at liquid nitrogen temperature (top) and in fixative after biopsy subdivision and warming (bottom). White zones indicative of ice formation are not visible before warming and did not appear during warming. **B.** No tissue cavities secondary to previous ice formation are observed, and no loss of ground substance is evident. Capillaries (plentiful near the top of the field) are intact and attached to adjacent neuropil. Two larger arterioles appear to show delamination of the tunica adventitia, whose outer layer remains attached to the surrounding neuropil (upper arrows) while inner layers and the arterioles themselves are contracted away from the outer layer. Two smaller arterioles including their outer tunica adventitia, are separate from the surrounding intact neuropil (lower arrows). Scale bar, 100 microns; 10X original magnification, H&E/Bielchowski’s dual staining to capture general ground substance. **C.** At higher magnification, cell shrinkage is apparent, with a rounded appearance of most cells, but tissue damage is not evident, and neural processes are visible as thin black filaments. Capillary separation from neuropil is not observed. Scale bar, 30 microns; 40X original magnification, H&E/Bielchowski’s dual staining.

**Fig.s 5B and C** display the larger scale histological state of human cerebral cortex after M22 perfusion, vitrification, and warming in M22 fixative at 4°C. Low power (10X, B) and higher power (40X, C) microscopy showed no evidence of loss of intracellular contents or other clear disruption of structural integrity beyond extreme dehydration. As observed in the rabbit brain, tissue cavities or other features suggestive of damage from freezing on cooling or devitrification upon rewarming are not observed, and capillaries appear intact and adjacent to neuropil. However, also as seen in the rabbit brain, arteriole separation from the neuropil (**A**, lower arrows) or from the outer layer of the tunica adventitia (**A**, upper arrows) can be observed (**A**), but the surrounding neuropil appears undisturbed. Also consistent with findings in M22-perfused rabbit brains, neural cells and processes are shrunken to the point of obscuring normally visible histological detail, with neurons appearing mostly as dark round objects (**C**).

The general impressions obtained from these light microscopic observations were confirmed by electron microscopy. **Fig. 6** displays the ultrastructural appearance of human cerebral cortical biopsies after M22 perfusion, vitrification in liquid nitrogen, and warming by placement in M22 fixative. At relatively low power (**A**, 1850X), the tissue is dense, obscuring detail. Many contorted shrinkage spaces can be seen adjacent to presumably osmotically active retracted features. Frank neuroanatomical discontinuities are not observed, and there is no indication of loss of ground substance, confirming the histological results of Fig. 5. The prominent black deposits near the center of the image (asterisk) are not identified. These are not observed in our rabbit brain studies, and are believed to be specific to this particular brain donor and to represent intracellular inclusions caused by an undiagnosed premortem degenerative condition or by an age-related process such as lipofuscin accumulation. Capillaries (C) are intact and adjacent to surrounding neuropil. At higher power (**B**, 11,770X), apparently osmotically shrunken axons (arrows) are visible within apparently intact myelin sheaths, replicating the results observed in rabbits (Fig. 4H). Fine neural processes are shrunken and separated by inter-process white shrinkage spaces but appear to be otherwise intact, with the exception of one process of reduced density (double asterisks). Synapses are somewhat difficult to identify at this magnification due in part to extreme tissue dehydration, but several putative examples are identifiable (circles). At still higher magnification (**C**: 22,420X; **D**: 24,110X), synapses are more unmistakable and are abundantly present (circles.) At these magnifications, shrinkage spaces help to demarcate what seem to be well-defined neural processes. Shrunken axons (asterisks, **C, D**) appear to be intact although separated from their surrounding myelin sheaths (denoted by double asterisks in **D**). Consistent with light microscopic observations, signs of frank damage secondary to ice formation are not observed.

**Fig. 6.**
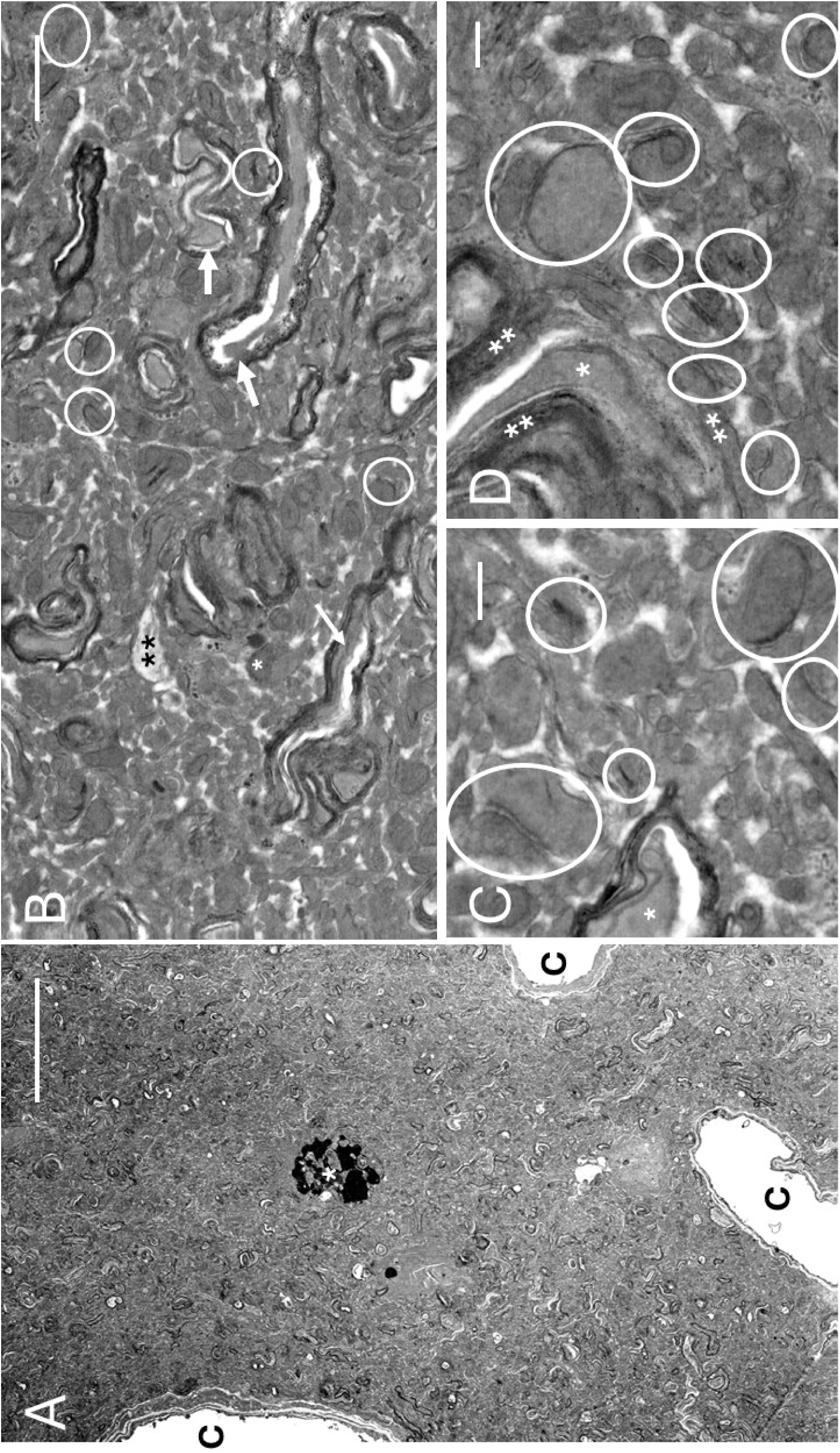
Ultrastructural appearance of human cerebral cortex treated as in Fig. 5B and C. **A.** Low-power (1850X) image, showing intact capillaries (C), dehydrated neuropil, and an inclusion (asterisk) thought to represent a pre-mortem pathology. Scale bar: 5 microns. **B.** At higher magnification (11,770X), shrunken axoplasm (arrows) is visible within largely intact myelin sheaths. A few likely synapses are circled. Numerous extracellular shrinkage spaces are visible. One area of abnormal cytoplasm is marked with double asterisks. Scale bar: 1 micron. **C.** Appearance at 22,420X. Synapses are clearly identifiable (circles). Shrunken axoplasm (asterisk) appears intact. Fine neural processes are separated by white extracellular spaces presumably caused by neural process osmotic shrinkage, implying the persistence of semi-permeable process membranes. Scale bar: 300 nm. **D.** Additional detail at 24,110X. Likely synapses are abundant (circles). Shrunken axoplasm (asterisk) appears intact within its surrounding myelin sheath (denoted by double asterisks). Scale bar: 200 nm.

### Brain Ultrastructure and Histology after Partial Brain Rehydration

To evaluate the reversibility of the structural distortions caused by dehydration, we investigated the effects of partial rehydration. Additional human cerebral cortical tissue samples were initially warmed to 4°C by placement in 75% M22 (**Fig. 7A**) or in 66% M22 (**Fig. 7B**) without fixative, held in either solution at 4°C with occasional agitation for 10-11 min, and then transferred to the same concentration of M22 in fixative. Again, no whitening of the tissue suggestive of devitrification was observed upon warming to 4°C. As shown in Fig. 7, a reduction of general staining density is apparent compared to Fig. 5, consistent with dilution of stainable molecules by partial tissue rehydration and re-expansion. There was also a progressive return of normal cortical pyramidal cell shape with increasing dilution, the rounded cell appearance shown in Fig. 5C being replaced by a restoration of spindle-shaped cells at 75% M22 (**A**) and a particularly striking reappearance of visually normal cell bodies and their apical and basal dendrite fields after dilution in 66% M22 (**B**). These changes suggest retention of cell membrane semi-permeability and osmotic responsiveness after warming from cryogenic temperatures. Capillaries remain intact and adjacent to surrounding neuropil.

**Fig. 7.**
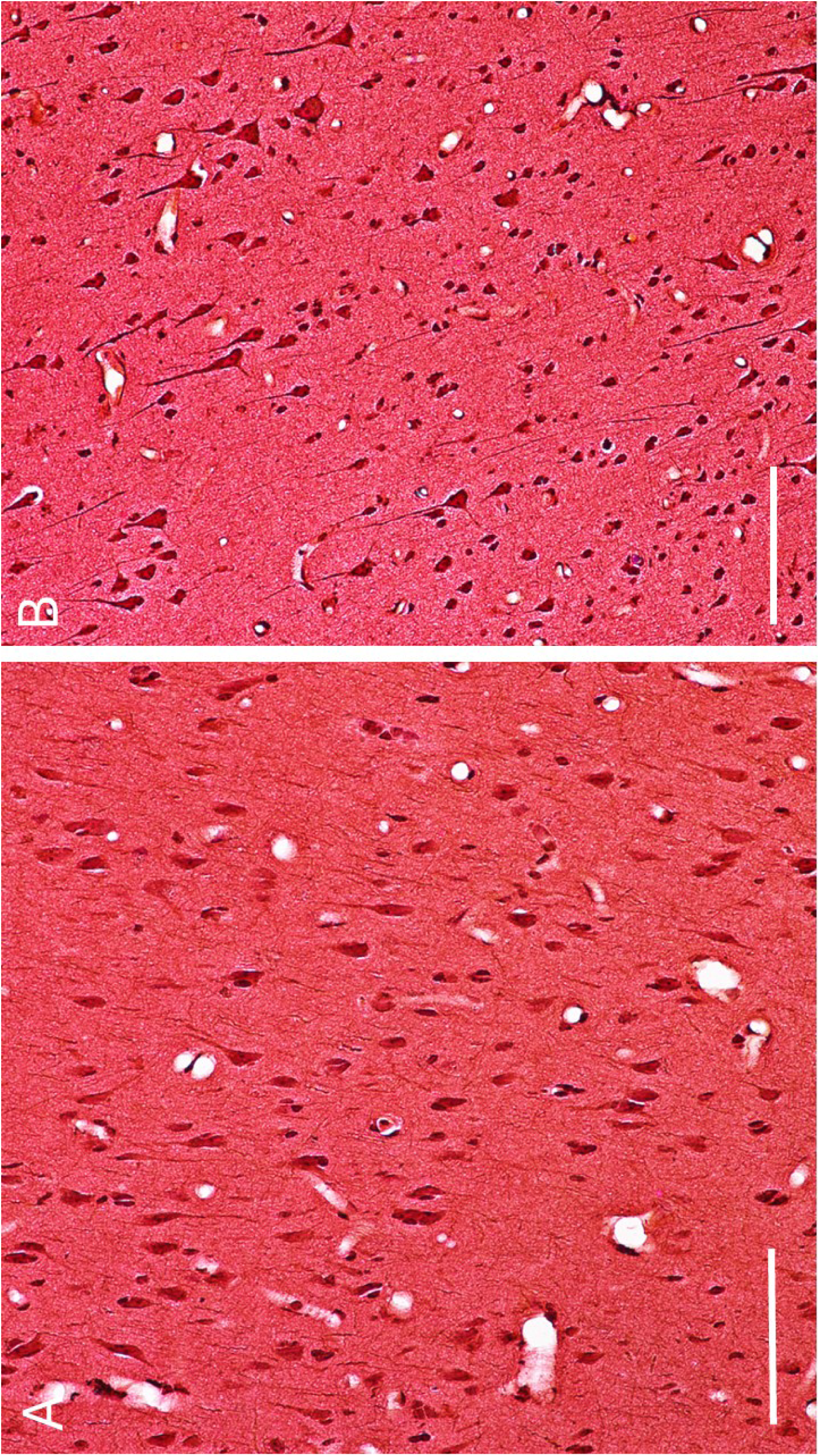
Osmotic responsiveness of vitrified/warmed human cerebral cortical neurons. **A.** Human cerebral cortical pyramidal cells after partial rehydration by immersion in 75% M22 (7.01M cryoprotectant) without fixative at 4°C, with swirling after 3, 5, and 8 min and fixation in 75% M22 fixative after 10 min followed by stepwise M22 washout. The staining intensity of the neuropil and cells is remarkably lighter, and there is a remarkable increase in visibility of pyramidal cell shapes and apical axons, compared to the appearance seen before rehydration (cf. Fig. 5B-C). Scale bar: 100 microns. **B.** Cerebral cortex after warming by immersion in 66% M22 (6.17M) without fixative at 4°C for 11 min followed by fixation in 66% M22 fixative and processing as in A. Tissue staining intensity is reduced compared to tissue before rehydration, and normal pyramidal cell shapes are strikingly restored, with well-visualized and abundant axonal projections. No loss of ground substance is apparent, and capillaries appear intact and attached to surrounding tissue. Some shrinkage spaces remain between pyramidal cells and the surrounding neuropil. Scale bar, 100 microns; 20X original magnification.

Parallel investigations were carried out on rabbit brains. Rabbit cephalons were first perfused with complete M22 for 30 min and then perfused with steadily decreasing concentrations followed by fixation. In the first series, concentration was slowly lowered to 5M M22, and the perfusate was then switched to 5M M22 fixative. Perfusion fixation was continued for 90 min at this concentration at around 10°C, after which the remaining M22 was then slowly removed by gradient perfusion dilution against 0M fixative overnight.

Dilution to 5M in this way resulted in remarkably little change in morphology in the intact brain compared to fixation in the presence of full concentration M22. **Fig. 8** displays remarkably shrunken CA1 cells (**A, B**), with prominent shrinkage spaces both extracellularly (ESS) and intracellularly (ISS), the latter being a phenomenon well documented to occur reversibly in cells shrunken by extracellular freezing without loss of viability [[31]]. As in the M22-perfused state, all anatomical features of the hippocampal cells and processes appear to be intact even though extremely dehydrated. **Fig. 8B** shows a larger shrunken myelinated fiber (SMF) near the point of origin of one CA1 cell apical axon (AX). This coincides in location with rare apparent gaps of the same size (apical gaps, or AG), whose presence is tentatively ascribed to artifactual loss of myelinated fiber contents during tissue processing. The affected fibers are shown at higher magnification in **Fig. 8C**. They appear to be fibers that pass through the stratum radiatum perpendicular to the apical axons. The lamellae of the myelin sheaths of these axons are partially separated, and the outer lamellae, while apparently intact, are sometimes separated from the surrounding entirely intact neuropil (arrows). The contained axons themselves (asterisks) are dehydrated but evidently intact. **Fig. 8D** shows, at higher magnification, abundant intact synapses (arrows), frequently crisp neuronal membranes, and sharp demarcations between cytoplasm and extracellular shrinkage spaces. The circular Fig. at the bottom right of Fig. 8D (double arrows) appears to consist of two apposed shrunken but intact myelinated fibers oriented perpendicular to the plane of the section. Mitochondria (M) appear to be intact. Although areas of possibly attenuated or pale cytoplasm can be seen (P), the predominant impression is one of intact neuronal membranes and cytoplasm.

**Fig. 8.**
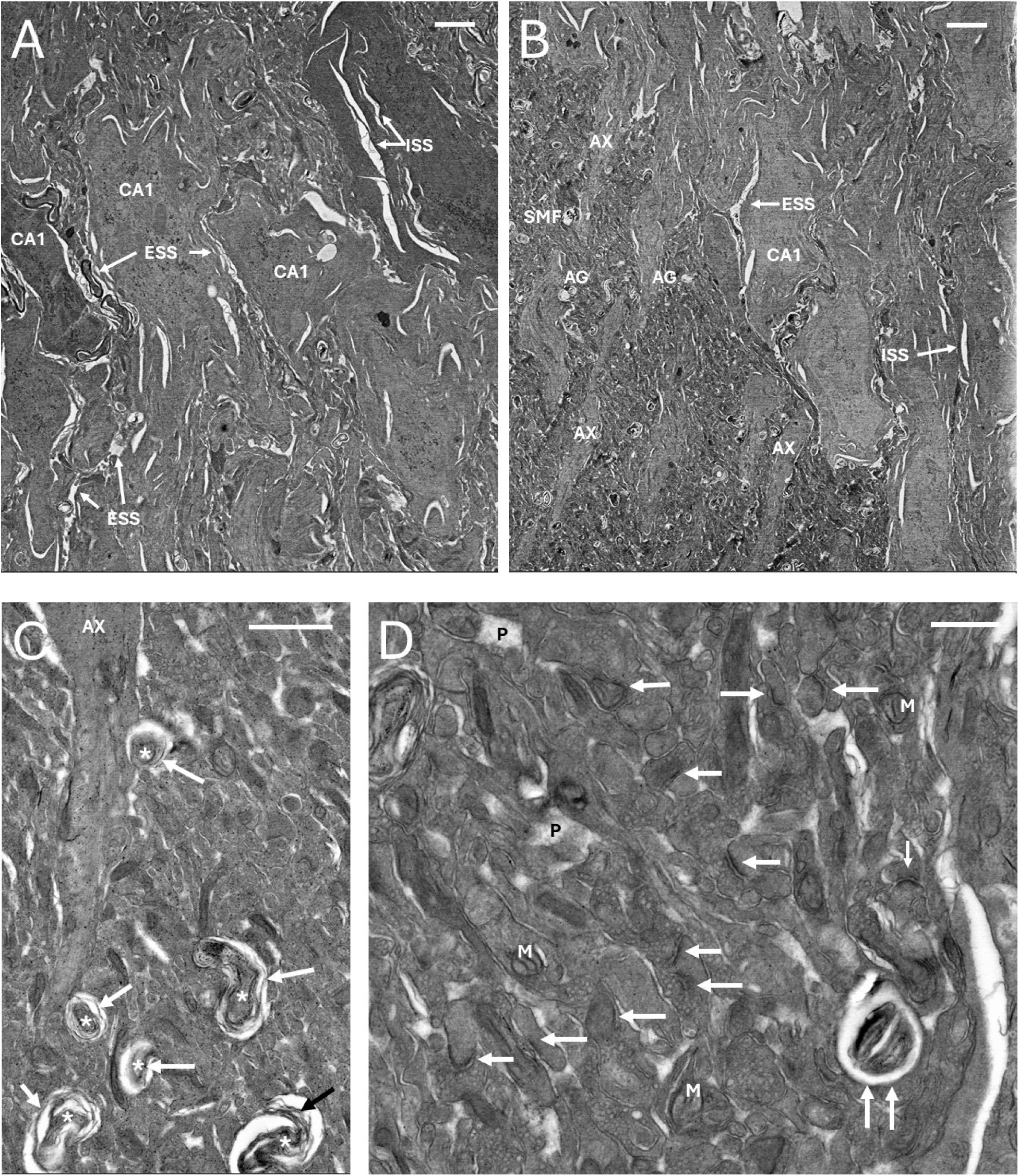
STEM images showing rabbit brain CA1 morphological variations following M22 perfusion followed by perfusion-dilution to 5M M22. **A.** Variably shrunken CA1 cells (CA1), with intracellular (ISS) and extracellular (ESS) shrinkage spaces still present. Apical axons are difficult to visualize due to tissue dehydration. **B.** Less severely dehydrated CA1 cells with visible apical axons (AX) and less pronounced shrinkage spaces. A shrunken myelinated fiber (SMF) and apical gaps (AG) believed to arise from SMF are visible near the points of origin of the apical axons. **C.** Shrunken myelinated nerve fibers passing through intact neuropil perpendicular to the plane of the section near a CA1 apical axon (AX). Arrows indicate the outer myelin layers. Asterisks: shrunken axoplasm. Numerous small extracellular shrinkage spaces are visible. **D.** Preservation of membrane and synaptic integrity in the SR despite persistent dehydration. Single arrows indicate intact synapses. Double arrows indicate a feature believed to be two apposed intact but shrunken myelinated fibers oriented perpendicular to the section. M: mitochondria. Extracellular shrinkage spaces frequently sharply display intact cell membranes. P: areas of what appears to be pale cytoplasm. Scale bars: 2, 3, 1, and 0.5 microns in A, B, C, and D, respectively.

To investigate the effects of further M22 dilution, rabbit brains were perfused with full M22 and then returned to 3M M22 and fixed as above. As shown in **Fig. 9**, this procedure resulted in dramatically less shrunken CA1 cells. Normal apical axons are seen in **Fig. 9A** and, at higher magnification, in **Fig. 9B**, but the surrounding neuropil continues to appear dark, and damaged myelinated fibers (DMF) remain present. The transverse myelinated fibers highlighted in **B** are more distorted than seen after dilution only to 5M M22 (Fig. 8C), and they still retain all expected anatomical features. In contrast to the DMF running perpendicular to the apical axons, one large myelinated fiber running parallel to the apical axons (panel **A**) contains a mildly shrunken but intact axon (arrow) and intact myelin. Deeper into the SR, neuropil examined under higher magnification (**Fig. 9C**) is less dehydrated than observed with dilution only to 5M M22 (compare to **Fig. 8D**), although some dehydration is still present. Nevertheless, rehydration is sufficient to visualize abundant intact synapses, and apparent presynaptic vesicles are visible (arrows). Shrinkage spaces are less prominent than at 5M M22, and some regions of attenuated or pale cytoplasm are visible (P), but most neuropil remains dark. In summary, Fig. 9 indicates that most anatomical features remain intact after M22 perfusion and that most ultrastructural distortion can be reversed by cryoprotectant dilution.

**Fig. 9.**
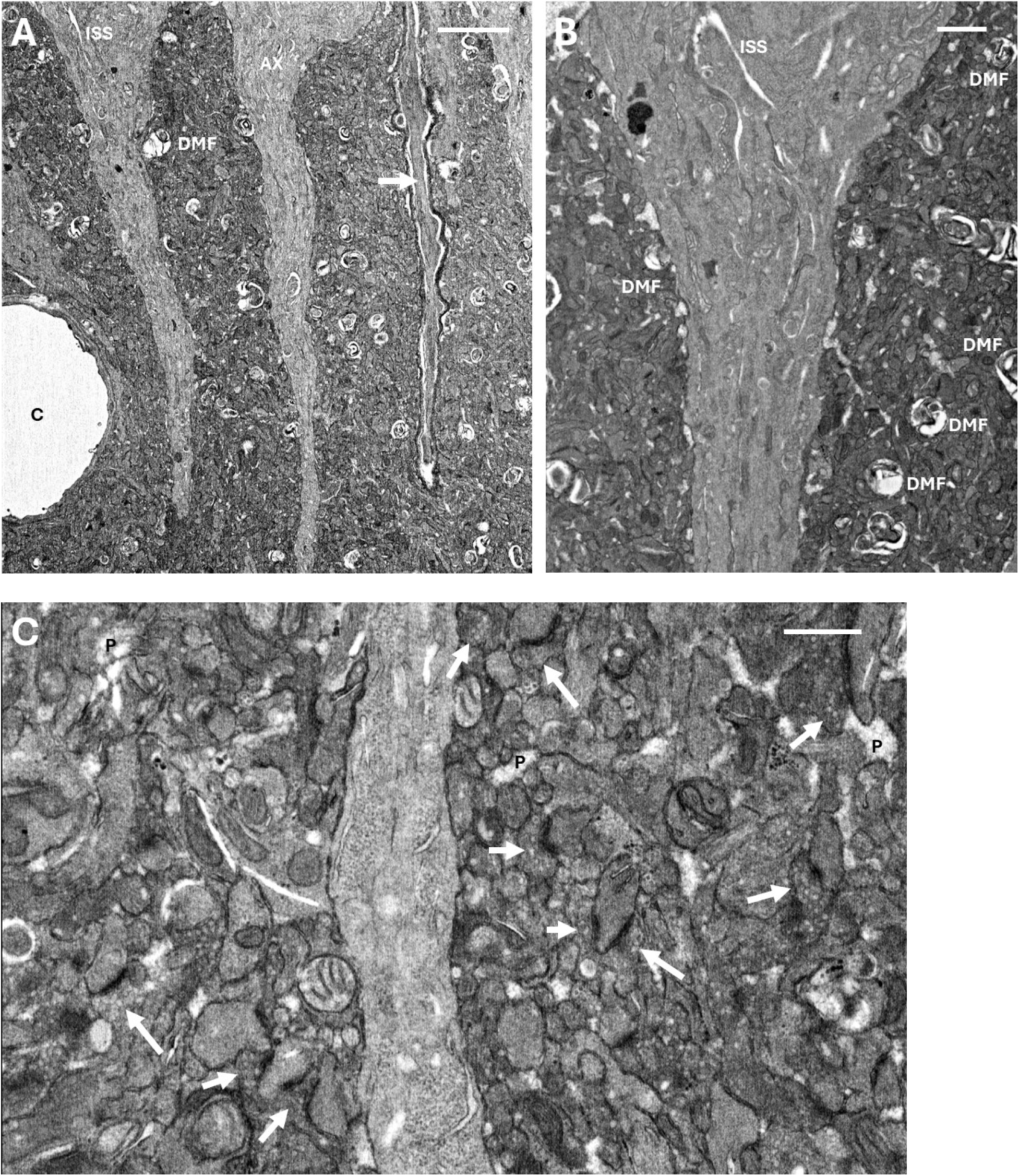
Return of less distorted morphology in rabbit hippocampus after perfusion with M22 followed by perfusion-dilution to 3M M22. **A.** Well-aligned CA1 cell apical axons (AX) appearing relatively normal except for minor intracellular shrinkage spaces (ISS). An identifiable damaged myelinated fiber (DMF) is visible, but a prominent myelinated nerve running parallel to the apical axons shows intact myelin and only mildly shrunken axoplasm (arrow). The capillary included in the field of view (c) is intact and adjacent to the surrounding neuropil. Scale bar: 3 microns. **B.** Slightly higher magnification, showing a variety of altered and putatively damaged myelinated fiber (DMF) morphologies. The apical axon appears intact other than for minor ISS, and the surrounding neuropil remains relatively dense. Scale bar: 1 micron. **C.** visibility of synapses and presynaptic vesicles (arrows) in the neuropil near the middle axon of A. Pale areas of cytoplasm (P) are also present. Scale bar: 500 nm.

One attempt to evaluate the effects of M22 dilution to 1M resulted in clearer images of presynaptic vesicles in more normal-density synapses (**Fig. 10A, B**). However, most neuronal structures appeared to be “exploded” (**Fig. 10C**). These results confirm that presynaptic vesicles are preserved in the presence of M22, but also show that the removal of more than two-thirds of perfused M22 requires measures to counteract the osmotic effect of retained intracellular M22, which is the presumed cause of the observed neuronal swelling and rupture. This may be approached using either slower M22 washout or employment of an extracellular osmolyte, or both. Studies of this kind were beyond the scope of the present investigation, but later work has confirmed that judicious use of osmolytes can prevent the disruption seen in Fig. 10 even when all M22 is washed out of the brain (Spindler et al., in preparation), thereby confirming that ultrastructural integrity is preserved in the maximally dehydrated brain.

**Fig. 10.**
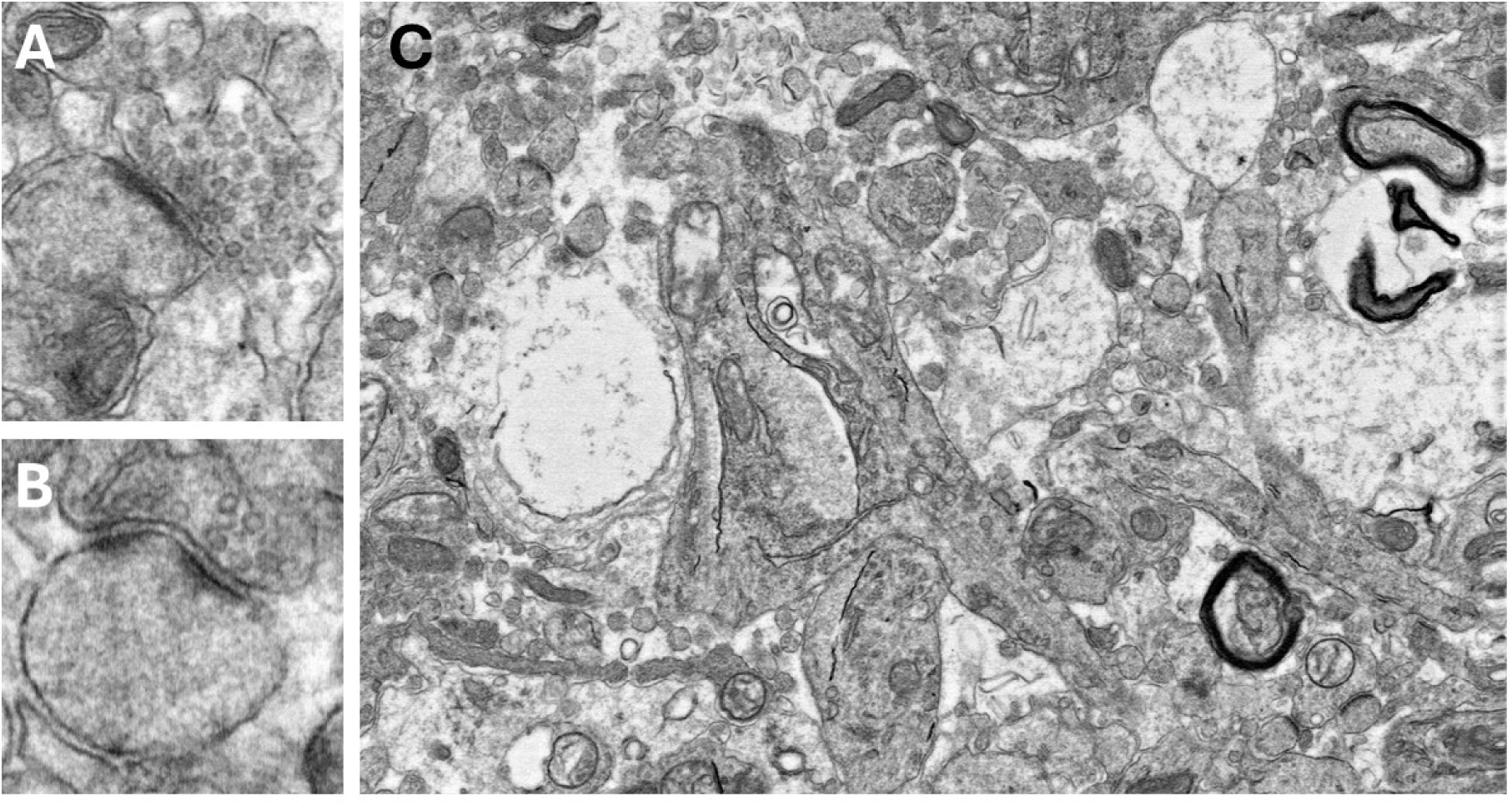
Effect of removing all but 1M M22. **A, B**. Synapses with well-defined presynaptic vesicles and postsynaptic densities. **C**. Ruptured neuronal compartments.

## Discussion

The studies reported here provide the first detailed survey and investigation of the ultrastructural and histological state of the brain and of regions taken from the whole brain after vitrification and rewarming and after the introduction and partial removal of vitrifiable concentrations of cryoprotective agents. The results confirm severe distortion of brain structure after the introduction of the M22 vitrification solution, but also the reversibility of brain cell shrinkage and much of the observed morphological distortion upon partial rehydration and indicate the retention of general membrane semipermeability and synaptic structural integrity. It seems apparent that vitrification is a far superior method of brain cryopreservation compared to freezing and that this method is sufficient to preserve almost all anatomical details and relationships in the brain without prior aldehyde fixation [[5]]. Most strikingly, our results provide the first evidence that human brains can, contrary to previous concerns [[5]], be perfused with a vitrification solution and vitrified hours after cardiac arrest with retention of histological and ultrastructural detail at least equal to what can be achieved in rabbit brains perfused and vitrified with no preceding period of warm and cold ischemia, raising new opportunities for scientific and medical applications of human and animal brain banking. However, our results also elucidate four primary types of structural injury that will require further analysis and mitigation in future studies.

The first and perhaps most fundamental of these is injury caused by the retained integrity of the blood-brain barrier (BBB) during cryoprotectant loading. The BBB is much more permeable to water than it is to cryoprotectants and therefore contributes significantly to brain dehydration and distortion. In unpublished studies (Spindler, in preparation), and in previous demonstrations of the efficacy of aldehyde-stabilized cryopreservation [[5]], we have found that using sodium dodecylsulfate (SDS) to open the BBB, as originally suggested by Pichugin (unpublished studies), can greatly mitigate or eliminate this source of injury. However, this method is hazardous, as it must be fine-tuned to permeabilize the BBB sufficiently to freely admit low molecular weight cryoprotectants but not higher molecular weight colloids, and may lead to entrapment of osmolytes and other undesired molecules in the extracellular space should the BBB reseal following CPA washout, which has been observed in some cases (Fahy, unpublished). Further, permeabilization of the BBB is not equivalent to permeabilization of neurons, glial cells, and other brain cells, and these may require additional study. However, complete or near-complete recovery of biochemical and neurophysiological properties is possible after vitrification and rewarming of brain slices and brain organoids [[14–16, 26, 33]], for which the BBB is not limiting, which implies that brain cell permeability to cryoprotectants is not a major problem. We also report here the first direct measurements showing the presence of substantial concentrations of perfused permeating CPA concentrations in brain tissue, showing that some CPA does cross the BBB and enter brain cells even without specific BBB opening.

The second type of injury is likely related to and may be a consequence of the first and consists of separation of arterioles from the surrounding brain parenchyma and/or delamination of some of the connective tissue layers of the tunica adventitia. Both were seen in both rabbit and human brain, whereas neither brain type exhibited capillary separation from surrounding brain tissue. Both phenomena may be caused by extreme osmotic vasodilation as water is drawn into the arteriolar lumen during CPA loading followed by arteriolar contraction as water fluxes slow and as intravascular pressure drops to zero as perfusion ends, potentially leaving a gap between the vessel wall and the surrounding dehydrated neuropil or between the layers of the tunica adventitia. The potential functional consequences of these changes, were the arterioles to be tested, are unclear. Damage to the tunica adventitia may weaken the ability of the vessel wall to withstand intravascular pressure or may otherwise increase arteriolar permeability, which could result in focal leakage. However, no signs of such effects were seen during washout of M22 as described. Opening the BBB might ameliorate such problems by accelerating CPA diffusion from the vascular lumen to the neuropil and thereby reducing osmotic vascular distention.

The third major type of morphological change induced by M22 perfusion is damage to myelinated nerve fibers. Cryoprotection of myelinated nerves was studied by Pribor and Nara [[34]] and by Menz [[35]]. The former authors found little effect of 15% v/v dimethyl sulfoxide on frog sciatic nerve electrical properties, but when the nerves were frozen, recovery of action potential amplitude was inversely proportional to the concentration of this agent, which they interpreted to mean that dimethyl sulfoxide did not penetrate the myelin sheath well. If correct, this would mean that at least frog myelinated nerves can withstand, in the absence of freezing, dehydration to an osmolality approximately 8 times higher than isotonic with little functional consequence, but no observations of the integrity of the myelin sheaths were made. Similarly, Menz found over 70% recovery of action potential amplitude in rat cutaneous nerves after abrupt or slow loading of 30% v/v dimethyl sulfoxide at 1°C, which implies tolerance of even more extreme osmotic dehydration by these myelinated mammalian nerve fibers, but again, the structure of the myelin was not examined. Kirschner and Caspar found that rabbit sciatic nerve myelin structure can be changed by dimethyl sulfoxide under extreme conditions, but that the changes are reversible when the CPA is removed [[36]]. We observe shrunken but apparently intact axoplasm within myelinated nerve fibers in both rabbit brains and human brain tissue and partial separation of the layers of the myelin sheath. The “apical gaps” noted in the hippocampus may represent peri-axonal shrinkage spaces that appear empty either because the plane of the section misses the enclosed axons or because the cross-sections of the enclosed shrunken axons have fallen out of the plane of the section during tissue processing due to their lack of connection to surrounding structures. Further definition of the nature of these gaps will require serial sectioning or FIB/SEM to trace their contours, but similar gaps and distortions can also be seen in non-cryoprotected control brains (data not shown), so we suspect these are artifacts. But even if these axons were to somehow be truly lost, the connectome would still be inferable because the myelin sheaths themselves remain to denote the paths of the axons.

The last kind of potentially significant injury is the presence of low-electron density areas that could be loosely termed “pale spaces” and that do not represent shrinkage spaces. These spaces are not obvious in the presence of peak CPA concentrations, but appear when the CPA is partially washed out. This suggests that these areas could be regions of preventable focal osmotic damage caused by local over-rehydration. Consistent with this possibility, we found that continuing dilution of M22 to 1M, while clearly demonstrating intact synapses and pre-synaptic vesicles, also led to areas of tissue disruption suggestive of excessive osmotic swelling. If this interpretation is correct, this injury should be preventable by using an osmolyte to counterbalance intracellular cryoprotectant osmolality during M22 washout or by removing M22 at a slower rate, or both, and ongoing studies are confirming this expectation (Spindler, in preparation). On the other hand, at peak concentration, some areas of seemingly coagulated cytoplasm can be seen, and if that effect does not quickly reverse itself upon dilution, it might manifest itself as pale areas. The significance and reversibility of both types of cytosolic change is unclear and merits further study, but pale areas are also frequently seen in control brains and in brains cryopreserved by ASC [[5]].

The results presented here are likely general in nature, but they are based on the use of just one CPA cocktail, M22, and one carrier solution, whose results may differ in detail from the results of using alterative vitrification solutions and carrier solutions. Further, in most of the studies reported here, the colloid support of the carrier solution (HES) was reduced to lower viscosity as M22 concentration was increased, and M22 was introduced in most experiments at-3°C, whereas M22 is normally introduced at-22°C in kidney perfusion experiments [[37]], and both of these factors could have affected our results in ways that are not presently clear. Clearly, the present results represent only a beginning, and much more research will be needed to fully define optimal methods for rendering brains vitrifiable and for restoring them to normal after rewarming or CPA washout. However, the demonstration of life support after vitrifying and transplanting rat [[38]] and rabbit [[39, 40]] kidneys and of EEG activity after high subzero temperature cryopreservation of brains [[2, 6]] suggests the feasibility of further progress.

Although imperfect, our results suggest the possibility of employing whole brain cryopreservation to preserve difficult-to-obtain brains for later pathological, neurochemical, ultrastructural, and even functional analysis [[1, 41]], including the brains of rare or endangered species, aged animals, or human brain donors with rare neurological conditions or diseases. Although it is necessary to add cryoprotectants to the whole brain by perfusion to preserve all brain regions simultaneously, it is not necessary to remove cryoprotectants by perfusion because brain subregions can be sampled for different studies and distributed to different laboratories at different times as is now commonly done by existing brain banks [[42]], and samples can be serially diluted at their final destinations to remove the cryoprotectants by diffusion [[5]], as was done for the human brain samples in the present study.

Finally, our observation that human brain cerebral cortical histology and ultrastructure was uniformly preserved despite two days of prior agonal hypotensive hypoxia and three hours of post-cardiac arrest hypothermic preservation before cryoprotectant perfusion provides the first direct evidence that human cryopreservation for medical time travel [[43]] may be feasible. This idea has captured much attention since it was first seriously introduced in 1964 [[44]], but to date, its scientific basis has been judged and debated mostly in the abstract and on the basis of missing or very indirect information and without direct inspection of human brain ultrastructure after preservation under practical conditions, an information gap that has finally now been at least partly filled. Although our present results describe the condition of only one human brain, they indicate less damage after vitrification than has been proposed to be reversible by methods of brain repair and resuscitation that have been forecasted through the use of advanced future technology [[45]]. Far more study is required, but our results with human cerebral cortical samples are consistent with results obtained with rabbit cerebral cortex and hippocampus, and cryogenic computed x-ray tomographic investigations have indicated that the entire human brain, and not just the cerebral cortex, can be successfully cooled to below the glass transition temperature without ice formation [[46]].

## Acknowledgements

We acknowledge the support of the late Dr. Harold T. Meryman, then at the American National Red Cross’s Blood Research Laboratory in Bethesda, MD (now closed), which made the experiments of Fig. 1 possible. We thank FEI, Inc., for preparing and making available Fig.s 1B and 1C and Dr. Alan Allenspach of the Department of Zoology at Miami University in Oxford, Ohio, for taking the photograph shown as Fig. 1D. We also thank Alicia Thompson, formerly the head of the Electron Microscopy and Microanalysis Center at USC (now retired), for taking the photographs shown in Fig. 3. The cat brain images of Fig. 2 were reproduced with the kind permission of the late Dr. Isamu Suda from the unpublished work of Suda, Kyoko Kito, and Chizuko Adachi. We thank Dr. Viktor Zach for removing and biopsying the human brain used for these studies. We are indebted to Linda Chamberlain for providing helpful supporting information and to Natalie Coles for facilitating and helping to enable the clinical research case described here. Nicholas Llewelyn kindly reviewed the manuscript and made helpful stylistic corrections. Supported by the Biomedical Research and Longevity Society, Inc., and by the Alcor Life Extension Foundation.

## Author contributions

G.M.F. conceived and directed most of the studies, participated in the animal studies, prepared the Fig.s, took the micrographs of Fig.s 2b and 4, and wrote the manuscript. R.S. and R.L. carried out confirmatory experiments, and R.S. reviewed and contributed to the manuscript. B.W. designed the DSC studies and participated in the design and conduct of the rabbit brain M22 washout experiments and edited the manuscript. V.V. carried out most of the rabbit brain experiments and devised and performed the analysis of tissue CPA concentrations. X.G. and A.S. performed most surgical procedures, and X.G. reviewed the manuscript. H.H. and S.G. carried out the human research procedures, and H.H. developed controlled cooling methodology for that study. B.T. and R.R. created most of the light and electron micrographs for this study, and R.R. reviewed the manuscript. S.B.H. provided medical support for the human brain study, and L.S.C. planned, arranged, and participated in the human brain study.

## Materials and Methods

### Animals and animal husbandry

New Zealand White rabbits were obtained from a rabbitry local to Bethesda, MD (for the studies of Fig. 1) or from Charles River Laboratories (for all other studies, which were carried out on specific pathogen free rabbits) and were housed according to USDA regulations and the Guide for the Care and Use of Laboratory Animals. All animal protocols were approved by Animal Care and Use Committees of the American National Red Cross (for the studies described in Fig. 1) and of 21^st^ Century Medicine, Inc. (for all other studies).

### Cryoprotectant, fixative, and carrier **s**olutions

The vitrification solution, M22, has been described elsewhere [[37]]. The carrier solution for M22 was LM5 [[37]] plus 1-5% w/v hydroxyethyl starch (HES) [mean molecular mass, 450 kD; degree of molar substitution, 0.7; obtained from Serumwerk, Bernburg, Germany, product HS# 3505 1050, or from B. Braun, Sabana Grande, Puerto Rico (now defunct), 460 kD, product X16-460-PR)] (HES protocols described below). M22 fixatives were prepared to contain 2.5% glutaraldehyde, 2% formaldehyde, 0.1M sodium cacodylate, and the concentrations of M22 solutes that were present in the non-fixative perfusate just prior to fixation. Glycerol for the studies of Fig. 1 was obtained from Sigma Chemical Company, St. Louis, MO. The glycerol perfusate contained 3.72M glycerol, 88.8 mM sucrose, 20 mM HEPES, 20 mM NaOH, 10 mM NaHCO_3_, 60 mM KCl, 5 mM reduced glutathione, 1 mM CaCl_2_, 1 mM MgCl_2_, and 5% w/v B. Braun HES. The blood washout solution used for most rabbit brain studies, KR8H, has been described elsewhere [[5]]. Blood was washed out with LM5 in the studies whose results are presented in Fig.s 3 and 4.

### Glycerol perfusion, freezing, and freeze-substitution

For the experiments described in Fig. 1 only, glycerol was perfused as described elsewhere [[1]]. Based on Smith’s observation that the hamster brain can tolerate conversion of 53-63% of its liquid volume into ice [[4]], it was desired to perfuse a concentration of glycerol that would prevent the ice content of the brain from exceeding 60% v/v at any temperature during freezing. This standard would require 40% of the initial liquid volume to remain liquid. Consequently, all solutes originally present, including glycerol, would be concentrated in the remaining liquid space by a factor of 1/0.4 = 2.5 fold. The highest concentration of glycerol that could be created in the spaces between ice crystals by freeze-concentration was found to be 73% w/w according to Rasmussen and Luyet [[47]], or 68% v/v, which is equivalent to 9.3M. Therefore, a rabbit brain was perfused with 9.3/2.5 = 3.72M glycerol in situ, removed, and cut into slabs that were frozen slowly to dry ice temperature. The frozen slabs were transferred into 50% ethanol/50% acetone containing 1% w/v osmium tetroxide as the fixative for freeze substitution [[12]]. The freeze-substitution medium was changed four times over the course of a week and then the brain pieces were warmed by placement in a-20°C freezer overnight and then placed into osmium-free medium at 4°C [[12]]. The tissue was then processed for light and electron microscopy according to standard methods [[12]].

### Rabbit brain M22 perfusion, vitrification and rewarming, dilution, and fixation

Initial cephalic blood washout and cooling were accomplished by sequential bilateral carotid perfusion to preclude cerebral ischemia as described [[5]]. After perfusion cooling to ∼4°C, the rabbit cephalons were perfused with LM5 containing 5% Serumwerk HES for 10 min at 3.5°C to establish an initial vascular resistance baseline. This solution was gradually transitioned to 5-8M M22 in LM5 over 80-105 min using a linear gradient against M22 containing 1% HES, and temperature was linearly lowered from 3.5°C to-3°C as concentrations increased from 2M to 5M. After a pause of ∼10 min, concentration was then switched in 1 or 2 steps to full concentration M22 in LM5 plus 1% HES at-3°C for 30-60 min as noted below. Specific examples of M22 perfusion protocols are given in Fig. SM-1.

Vitrification was accomplished by exposing rabbit cephalons to rapidly-moving liquid nitrogen vapor in a modified Linde BF1 forced-convection cooling chamber (no longer manufactured). Vapor temperature was typically set to-100°C for 30 min, to-120° for another 30 min, and then to-130°C until the deep pharyngeal temperature reached-123°C or below, which typically required a total cooling time of ∼90 min. Rewarming was carried out in the BF1 unit with the assistance of four specially-installed 500-watt heating elements. The atmosphere in the BF1, which was rapidly stirred by a centrifugal fan located at the bottom of the chamber during both cooling and warming, was raised to ∼0°C over 10 min and held constant until the deep pharyngeal temperature reached ∼-20°C, which typically required a total warming period of around 60 min.

Graded M22 washout prior to fixation was done linearly at a rate of approximately 50-70 mM/min as described in the Supplemental Materials while perfusion temperature was gradually raised toward 3.5°C as described elsewhere [[37]]; see also **S1 Fig**. When the target concentration (5, 3, or 1M) was reached, washout was halted, and perfusion was switched to the same M22 concentration in the above-described fixative for 90 min. The remaining M22 was then generally removed overnight (over approximately 15-23 hours) using a linear gradient against the same base fixative solution minus M22 components. Brains fixed in the presence of complete M22 were removed, dissected, and diluted stepwise over ≥∼2 weeks by immersion in successive M22/fixative solutions each containing 33% less M22 than the previous solution until the final cryoprotectant concentration was <150 mM. Each dilution step lasted at least 1 day. The slabs were then transferred to 0 mM M22 fixative.

### Human brain M22 perfusion and cryopreservation

A 73 year-old terminal male pancreatic cancer patient provided informed consent and completed legal arrangements in November of 2014 to donate his brain for scientific research to enable the studies reported here. He was admitted to a hospice, where he was given pain medications (morphine and Ativan) and then became anorexic. Four days later, he became unresponsive and hypotensive (93/73 mmHg), with an oxygen saturation of 88%. The next day, his blood pressure was 78/56 to 81/51, with an oxygen saturation of 87%, and SpO2 was maintained at 82-89% with pressures of 88/56 on the following day, when a fentanyl patch was added. Blood pressure thereafter declined to 79/44 to 74/41, SpO2 declined to 81%, and septicemia was noted. The following morning, blood pressures descended to 47/23, with a respiratory rate of 8 and an SpO2 of 75%, and death was pronounced 19 min later, at 9:50 am, December 3^rd^, 2014. The subsequent details of processing for cryopreservation are described in the Supplemental Materials.

### Light and electron microscopy

Samples from whole brains were fixed and washed free of M22 as described above. For electron microscopy of fixed rabbit brain samples after M22 washout, a Stadie-Riggs microtome blade (cat. No. 6727C18, Thomas Scientific) was used to prepare a roughly 1 mm-thick whole-brain coronal section that transected the region of the hippocampus dorsal to the approximate center of the thalamus. Rectangular samples including the most dorsal aspect of the CA1 band and the overlying cerebral cortex were prepared with a scalpel blade (sampled area denoted in Fig. 3A). From these macro samples, smaller (<1 mm^3^) sample pieces were prepared from both the cortex and the portion of the hippocampus containing at least the CA-1 cell band for further processing. Human brain cerebral cortical biopsies obtained from both the right and left dorsal and ventral frontal lobes were initially transferred directly into liquid nitrogen for later analysis. Four years later, they were divided into ∼0.3-4 mm^3^ slab-shaped pieces (maximum length, width, and height dimensions: 5, 1.5, and 1.4 mm, respectively) under liquid nitrogen and transferred to M22 fixative at 4°C or at room temperature and swirled immediately to rewarm them and again about 5 and 30 min later to eliminate unstirred solution layers around the tissue. After overnight storage at 4°C, 10 cryoprotectant washout steps were performed over 35 days, each step lowering the cryoprotectant concentration by 33.3%. On day 36, having reduced the concentration to 9.45*0.6667^10^ = 0.164M, the tissue was transferred to cryoprotective agent-free fixative. All human biopsies intended for light microscopy were shipped to American Histolabs (Gaithersburg, MD; now defunct) for paraffin embedding, sectioning, and staining with a combination of hematoxylin-eosin and Bielchowski’s stain. Samples intended for STEM examination were cut to a size of ∼1 mm^3^ and processed as for rabbit brain samples. Samples fixed after previous dilution were processed as described in the Results.

After cryoprotectant removal, fixative was removed by holding samples overnight in a refrigerator at 4°C in small vials containing 2 ml of 0.1M cacodylate buffer without fixative. The samples were washed the following morning with two more changes of 4°C buffer lasting 20 min each. The buffer was then replaced with a secondary fixative consisting of cacodylate-buffered 1% w/v osmium tetroxide (OsO_4_) and 1.5% w/v potassium ferricyanide (added as an additional membrane contrast agent) for 1 hour, after which the samples were transferred to the same solution without ferricyanide for another hour. The OsO_4_ was washed out using two 10 min washes at room temperature (RT) in MilliQ water, and the water was replaced with 1% w/v uranyl acetate in water for 1 hour. Subsequent dehydration, embedding, and sectioning procedures for samples intended for STEM were routine and are described in the Supplementary Materials. Scanning electron microscopy of brain slabs was carried out at USC by A.T. after whole brain perfusion-fixation and perfusion-unloading of CPA as described above, brain slab preparation by hand-cutting with a Stadie-Riggs microtome blade, and critical point drying with pressurized CO_2_ followed by coating and scanning according to standard techniques.

### Differential scanning calorimetry

Samples intended for HPLC and DSC studies were warmed rapidly by being dropped into room temperature M22, which prevented frost deposition on the sample and surface evaporation, and then immediately blotted to prevent uptake of bath M22 into the samples. No whitening of the samples was noted during rapid warming. Two DSC protocols were used to determine ice formation tendency, one to detect ice formation during slow cooling (failure to fully vitrify), and one to purposefully form clearly detectable amounts of ice so that the concentration of cryoprotectant in the tissue could be estimated from the observed ice melting point. In the first protocol (Protocol 1), samples were cooled from 0°C to-90°C at 1°C/min and then warmed to +30°C at 160°C/min to detect ice formed on cooling by the presence of a melting peak on warming. In the second protocol (Protocol 2), samples were deliberately placed at risk of nucleation by cooling at 100°C/min to-120°C (at which temperature ice nucleation rates are maximized [[32, 48]]), holding for 3 min at that temperature, warming to-60°C and holding at that temperature for 10 min to allow developed nuclei to grow to significant size, recooling to-80°C at 100°C/min and holding at-80°C for 1 min (to avoid startup artifacts upon subsequent warming), and then warming at 5°C/min to +10°C to measure the melting temperature of previously formed ice. All experiments were conducted using a Perkin-Elmer DSC-7 and Perkin-Elmer 0219-0062 aluminum sample pans with tissue sample masses between 10 and 14 mg.

### HPLC determination of cerebral tissue cryoprotectant concentrations

Rewarmed and immediately blotted brain tissue samples were placed into 2 ml of a space marker solution consisting of 2.6% w/v glycerol and 2.844% w/v trichloroacetic acid in water and contained in a high precision (mean measured accuracy, ±0.25% at 2 ml) 7 ml graduated cylinder (shortened from 10 ml to permit subsequent homogenization in the cylinder). Tissue volume was determined as the volume increment in the cylinder, as determined visually and by measuring displacements less than 0.1 ml to the nearest 1 µl by aspiration with a P100 pipet, with return of aspirated fluid to the cylinder. The sample was homogenized with a Fisher Scientific TissueMiser homogenizer, and 1.5 ml of the homogenate was transferred to a 2 ml Eppendorf microcentrifuge tube and spun at 16,000 x g for 5 min. 100 µl of supernatant was diluted with 900 µl of MilliQ distilled/deionized water, and the mixture was filtered through a Whatman Spartan 0.22 micron, 13 mm diameter syringe filter (regenerated cellulose). A 20 µl HPLC sample loop was flushed with either 100 µl of sample or with 100 µl of 10X-diluted space marker solution as a control, and the loop contents were then injected through a pre-column 8 mm ID Shimadzu SecurityGuard (KJO-4282) guard cartridge with an internal 3.2 mm ID Carbo-H column (AJO-4490). The column was a Phenomenex 8µ Rezex^TM^ Organic Acid H+ LC column (00H-0138-KO; 300 x 7.8 mm), and separation was done using isocratic 5 mM H_2_SO_4_ as the mobile phase for 55 min at 35°C with a backpressure of approximately 470-530 PSI. RI peak areas were calibrated against separate M22 and glycerol dilution standards ranging from 25-200 fold dilutions of M22 and 6.5-52-fold dilutions of space marker glycerol to determine molar concentrations of M22 components and glycerol in the injected samples using separate calibration slopes and intercepts for each chemical. A 100 µl tissue sample would experience a dilution of approximately (2.1/0.1)x(10) = 210-fold, which is at the limit of the calibrated concentration range. By mass conservation, C_CT_ = 10C_CH_(V_H_+V_L_)/V_L_, where C_CT_ is the concentration of a given cryoprotectant in the tissue liquid space before homogenization, C_CH_ is the concentration of the same cryoprotectant in the homogenate (multiplied by 10 to correct for the 10-fold dilution of the homogenate prior to HPLC analysis), V_H_ is the volume of homogenization solution (2 ml), and V_L_ is the liquid space within the tissue. V_L_ is calculated from the dilution of space marker glycerol by the tissue as V_L_ = V_H_(C_Gb_/C_Ga_-1), where C_Gb_ and C_Ga_ are the glycerol concentrations before and after tissue addition, respectively. The liquid fraction of the tissue is V_L_/V_T_, where V_T_ is the total tissue volume, equal to the volume increment measured in the graduated cylinder after addition of the tissue. For purposes of presentation, C_CT_ is normalized to C_CM_, which is the concentration of the cryoprotectant in M22. Alternatively, C_CT_/C_CM_ can be equated to the apparent cryoprotectant space relative to the glycerol-accessible tissue space.

## Supplementary Materials

### Rabbit Brain M22 perfusion, vitrification, and CPA washout methods

The protocols used to generate the results presented in Fig.s 4 and 8-10 are described in Fig. SM-1 A, B, C, and D, respectively, below.

**Fig. SM-1.**
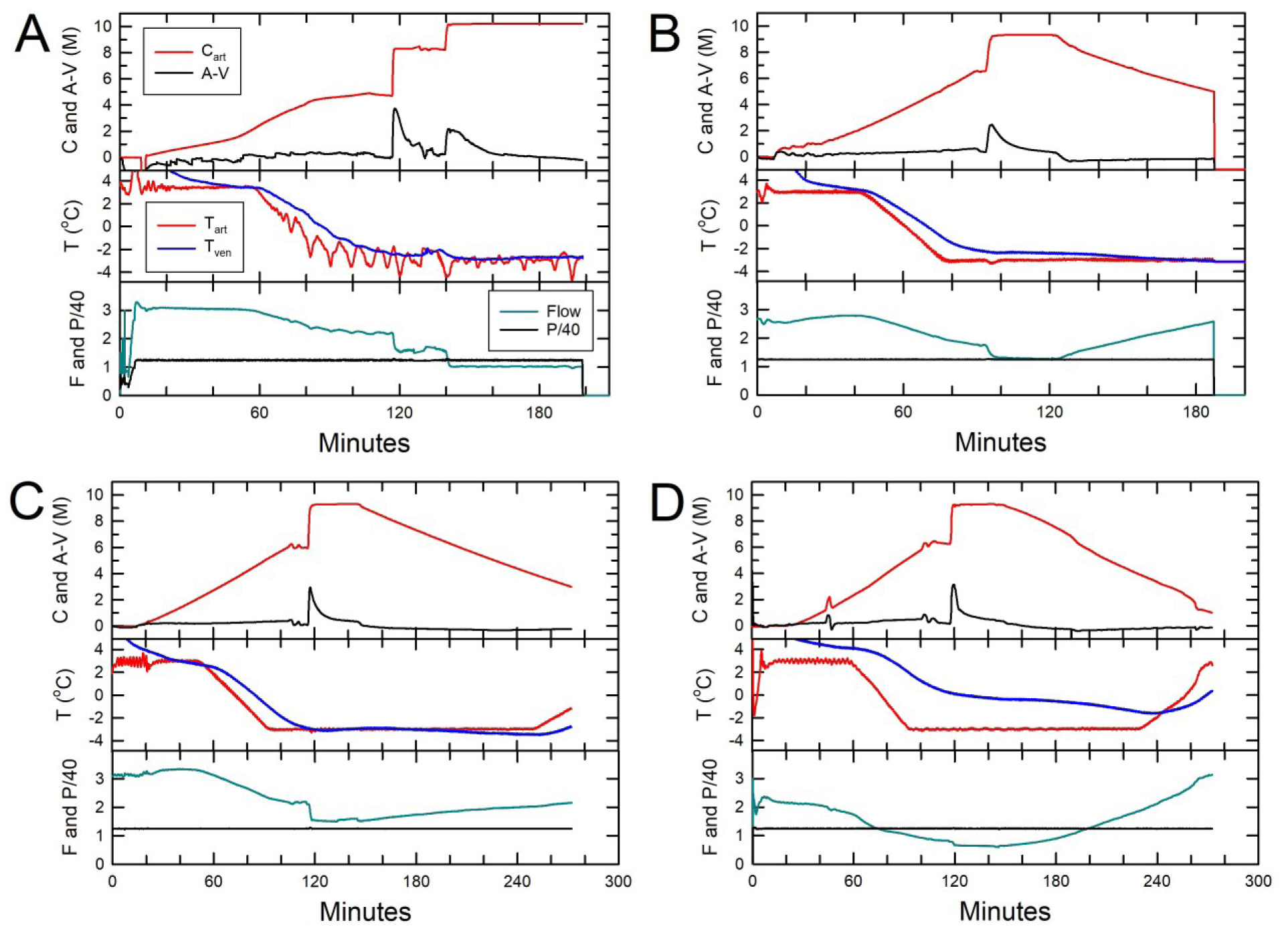
C (red) = molar concentration; A-V (black) = arteriovenous molar concentration difference; T (°C) = temperature: red = arterial, blue = venous; F (dark cyan) = flow [(ml/min)/10g], P/40 = pressure (mmHg)/40.

### Routine sample processing for scanning-transmission electron microscopy (STEM)

After post-fixation with uranyl acetate as described, samples were rinsed for 15 min twice in water and were dehydrated in 12 min steps at room temperature in 50% v/v, 70% v/v, and 95% v/v ethanol followed by three more 12 min dehydration steps in 100% ethanol. The samples were then transferred through three 15 min washes in 100% propylene oxide (PO) and placed in a 1:1 mixture of PO and Embed-812 epoxy resin for 1 hour followed by a 1:2 mixture of PO and resin at RT on a rotator overnight. The samples were finally placed into pure resin the following morning for at least 3 hours on a rotator to complete infiltration and then placed into individual, labeled molds filled with resin and polymerized for two days in a 65°C oven. The hardened blocks of polymerized resin were hand-trimmed and cut with a glass knife to make ∼0.7µm-thick sections for toluidine blue staining for light microscopy. Selected areas were then further cut with a diamond knife into ∼70 nm sections for STEM imaging. Sections were coated with ∼10 nm of evaporated carbon to enhance stability and examined on 200 mesh copper grids in a Zeiss Supra 40 scanning electron microscope at 28 kV using a STEM detector.

### Human cerebral cortical sample dilution schedule

The procedure for removing M22 from cortical biopsies is described in Fig. SM-2.

**Fig. SM-2.**
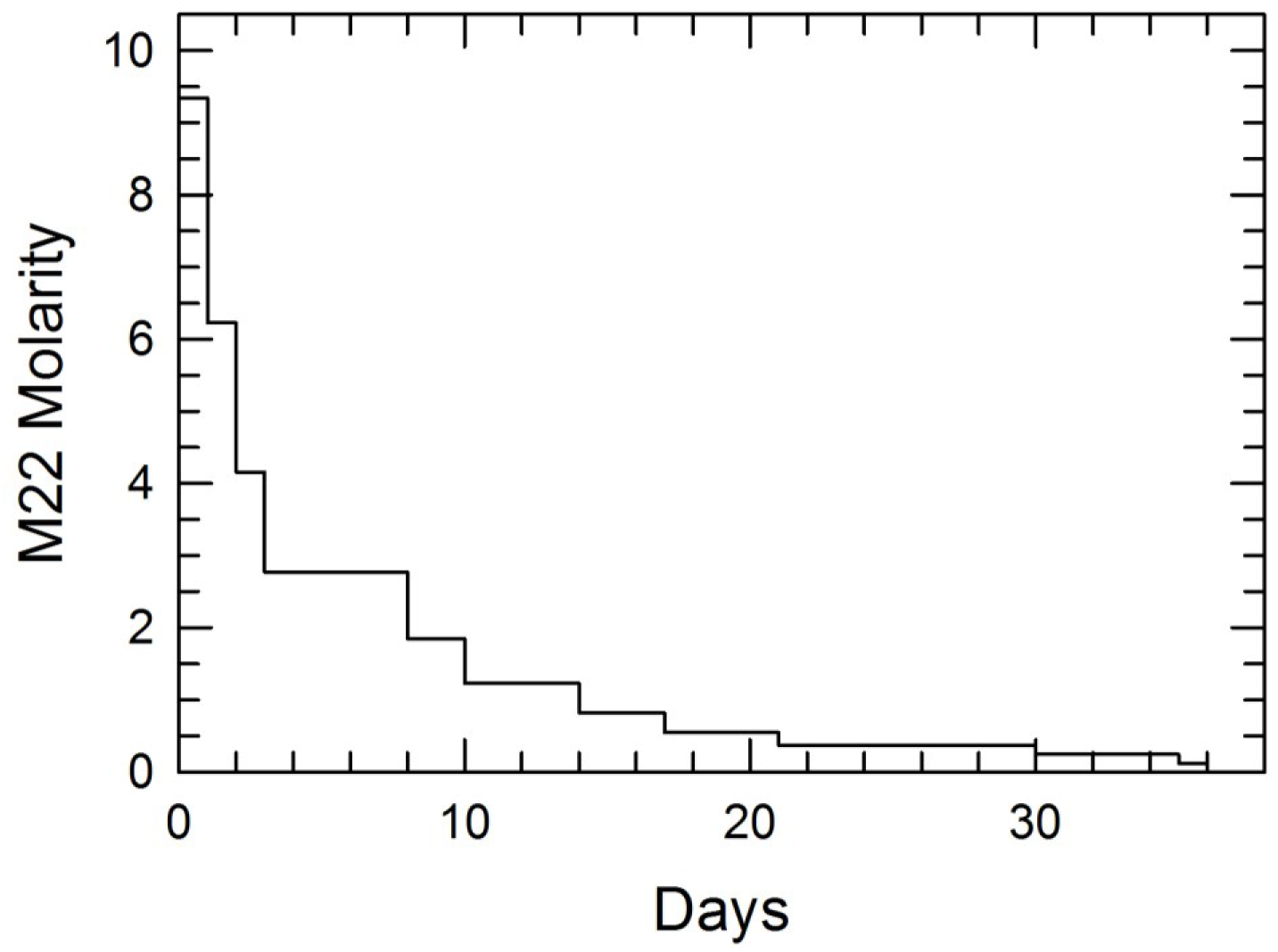

### Human Cerebral Cortical M22 Uptake

Table SM-1 shows preliminary data on the distribution of M22 components within the glycerol-accessible liquid space of the cerebral cortex. The liquid volume fraction of the tissue (estimated as the fraction of total tissue volume accessible to glycerol) was calculated to be 55%. Assuming a normal liquid volume fraction of 80% [[49]], approximately 31% of brain liquid was subtracted by dehydration. Given an isotonic cytoplasmic intracellular protein concentration of 15% w/w (higher in organelles) [[49]], dehydration would have concentrated intracellular proteins to about 22%, which is expected to contribute substantially to stability against ice formation [[50]]. The carbohydrates used as components of the carrier solution for M22 (LM5) had access to approximately 50% of the available liquid space, which likely reflects their inability to penetrate cell membranes and may be influenced by incomplete permeation through the blood-brain barrier. Nominally cell-permeating cryoprotectants were distributed throughout a greater fraction of the glycerol space. However, even when unable to distribute fully into those spaces, they must induce sufficient dehydration to raise the concentrations of other intracellular solutes sufficiently to approximately equate intra-and extracellular osmolality, thus increasing intracellular resistance to ice formation indirectly.

**Table SM-1:** Apparent Volumes of Distribution (VD) of M22 Perfusate Constituents in Human Cerebral Cortex.

| Table SM-1: Apparent Volumes of Distribution (VD) of M22<br>Perfusate Constituents in Human Cerebral Cortex<br>(As a Percentage of the Glycerol-Accessible Space) |  |
| --- | --- |
| Perfusate Constituent | VD (%) |
| N-methyl formamide | 103.8 |
| Ethylene glycol | 102.6 |
| 3-methoxy-1,2-propanediol | 81.1 |
| Dimethyl sulfoxide | 79.2 |
| Formamide | 54.3 |
| Glucose | 49.2 |
| Lactose | 49.3 |
| Mannitol | 46.7 |

### Human cerebral cortical stability against ice formation on cooling and warming

Fig. SM-3 shows representative DSC warming profiles of M22-perfused human cerebral cortical samples (n=3) after previous cooling to-90°C or below (see Materials and Methods for details). **A.** Warming at 160°C/min after cooling to-90°C at 1°C/min. At this warming rate, any ice that formed during cooling would be detected as a large melting peak between-55°C and 0°C; no such peak is observed. **B.** Warming at 5°C/min after intentionally inducing ice nucleation at-120°C and allowing ice growth at-60°C so as to facilitate detection and measurement of a tissue ice melting point on subsequent warming. As in A, any ice formed should be detectable as a distinct melting peak between-55 and 0°C in the highly amplified scale of this image. No ice was seen in any sample subjected to this procedure. After heat flow stabilization, deviations from thermogram linearity are due to imperfections in the instrument baseline at this amplified scale, not thermal events in the sample.

**Fig. SM-3.**
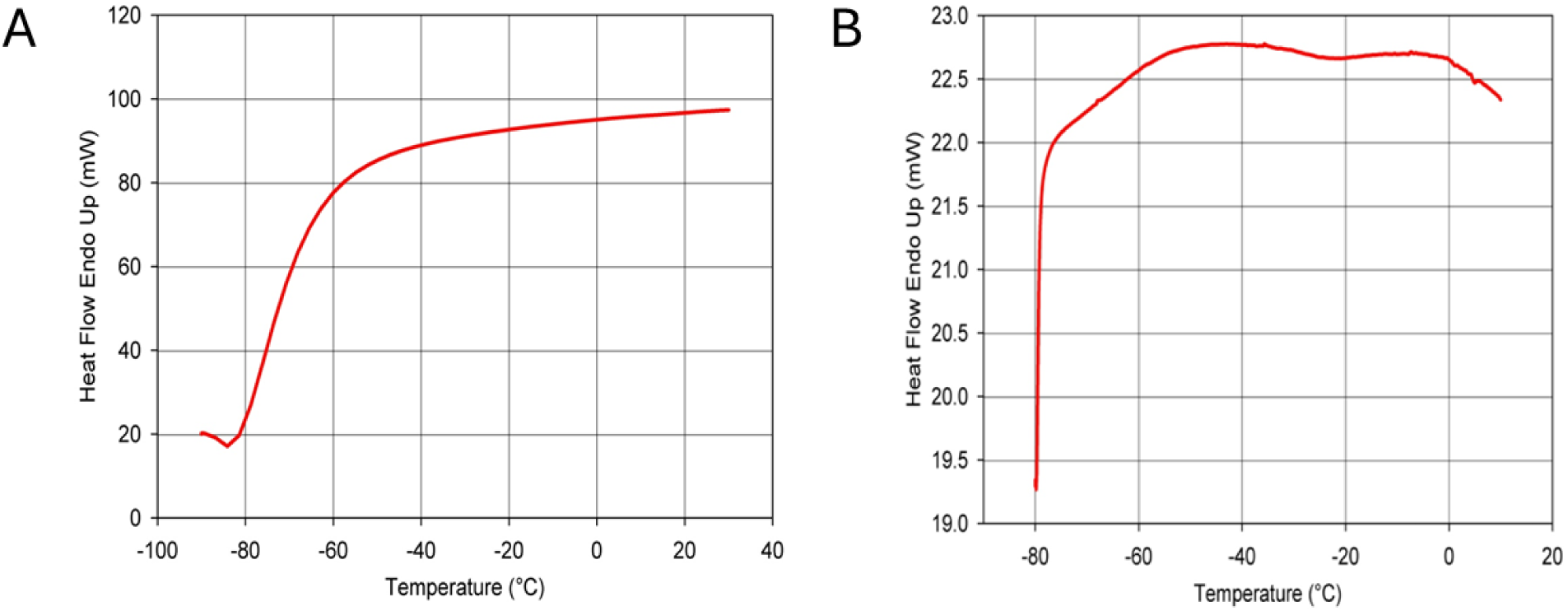

### Donor brain preservation

The donor was placed into an ice bath 1 min after the legal declaration of death based on cardiorespiratory arrest (minute 1) and received intravenous propofol (200 mg in 20 ml), filter-sterilized streptokinase (250,000 units), and acetylsalicylic acid (300 mg in 10 ml of 1M THAM) after 7-8 min of cooling. The medications were circulated using a Lucas 2 resuscitator (Physio-Control, Inc., Lund, Sweden) at >100 compressions/minute beginning 2 minutes later. Sodium citrate (50 ml of 20% w/v) was administered IV 4-5 min after initiating circulatory support, and 100,000 IU of heparin was given immediately thereafter, in two 50,000 IU boluses. At this point, at minute 16, or 15 min after the onset of ice bath cooling, tympanic membrane temperatures were 26.1-34.2 °C. An airway was installed at minute 17, and 4-hydroxytempo (5 g in normal saline, filter-sterilized) was given and infusion of an epinephrine-vasopressin mix (30 ml, 1:1,000 epinephrine and 100,000 IU vasopressin) was started within the next two min. Mechanical ventilator support began at minute 21, and an additional 25 ml of 1M THAM was given at minute 22 followed by 5-methylisothiourea (400 mg), gentamicin, ketorolac, another 25 ml of THAM, and 50 ml of 25% mannitol by minute 30. Transportation from the hospice began at minute 35, with tympanic temperature readings of 18 and 32°C. A proprietary emulsion containing melatonin, vitamin E (d-alpha tocopherol), alpha phenyl t-butyl nitrone (PBN), and carprofen in Cremophor EL (Vital-Oxy) was administered over minutes 70-84 and the donor arrived at the laboratory at minute 84 with tympanic readings of 16.4 and 21.5°C.

After surgical preparation for bilateral carotid perfusion, selective cephalic perfusion with LM5 containing 5% HES began at 12:31 pm (minute 161). Perfusion was initiated at a nasopharyngeal temperature of 17.4°C, an arterial temperature of 5.9°C, a right jugular temperature of 8.9°C, an environmental temperature of 16°C, and an arterial pressure of 130 mmHg. Perfusion with M22 plus 4% HES began 16 min later (at minute 177). Concentration was raised to ∼5M over the course of 91 min (mean ramp rate, 55 mM/min), reaching this first plateau concentration at minute 268. During this gradual M22 introduction phase, the perfusion pressure remained constant, arterial and venous temperatures descended to and remained at 4°C, the environmental temperature was quickly set to 3°C, and the nasopharyngeal temperature remained between 15 and 20°C. Upon reaching this first M22 concentration plateau, the environmental temperature was quickly reset to-4°C, resulting in a nasopharyngeal temperature of 7°C, the arterial pressure was reset to 100 mmHg, and the arterial temperature was gradually lowered to-3°C while the M22 concentration was held approximately constant for an additional 33 min to promote cryoprotectant distribution into the tissues. After this first plateau phase, the arterial concentration was increased to the target of 105% of the full concentration of M22 (∼9.9M) within about 22 min and maintained between 102 and 105% of full M22 for an additional 53 min for final equilibration while the other perfusion parameters remained constant. The perfusion ended at minute 376, or 6 hours and 16 min following the legal declaration of death.

After a one hour and 8 min delay, the brain was surgically removed and placed on a Teflon film supported by a cushion of Dacron wool within a metal bowl, and bilateral dorsal and ventral cortical biopsies were collected and dropped directly into liquid nitrogen in small sample containers. These procedures required 29 min to complete. The brain was then cooled by lowering the bowl to a fixed position within an MVE TA-60 dewar precooled to-100°C with cold, dry nitrogen gas. The bowl was covered with a sheet of aluminum to protect the brain from any stray droplets of liquid nitrogen, and the dewar was closed with an insulated lid equipped with a liquid nitrogen injector and a fan to stir the gas space within the dewar. The injector nozzle was separated from the sample by a metal barrier attached to the insulated lid. Temperature within the dewar was controlled with a LabVIEW program that operated an external liquid nitrogen solenoid valve inlet. The program set the initial temperature to-100°C and changed temperature thereafter at linear rates of ≤1°C/hour between different target temperatures. Cooling proceeded initially from-100°C to-147°C over 48 hours, after which temperature was returned at a linear rate to-140°C over an additional 102 hours. After storage for an additional 13.5 days, the brain was transferred under isothermal conditions to a permanent-140°C storage unit [[51]]. Before entering long term storage, but within a thermally controlled environment, the brain was inspected and photographed to document the presence or absence of surface cracks. Surface cracks were not observed. Extensive research has indicated that fracturing in large organs caused by cooling to below the glass transition temperature is always visible on the organ surface, internal cracks being absent when external cracks are not present (Fahy et al., unpublished observations).

## Notes

### Competing Interest Statement

The authors have declared no competing interest.

### Summary of Updates

Dr. Harris' first name is spelled "Steven" rather than "Stephen" as originally submitted. The sole purpose of this revision is to correct the spelling error in his name.

